# UNC5D Promotes Retinoblastoma Progression Through Association with NTN1 and Activation of PI3K/AKT Signaling

**DOI:** 10.64898/2026.11.23.768624

**Authors:** Aditi Karmakar, Subhankar Bose, Shubhrajit Barman, Amit Kumar Srivastava, Senthil Kumar Ganesan

## Abstract

Retinoblastoma (Rb) is the most common pediatric intraocular malignancy, yet the molecular mechanisms that mediate tumor progression beyond RB1 loss remain incompletely understood. Through our previous integrative multi-omics analysis, UNC5D was identified as a candidate regulator of retinoblastoma pathogenesis. In the present study, we investigated the functional role and molecular mechanism of UNC5D in Rb. We found that UNC5D expression was significantly elevated in Y79 and Weri-Rb1 retinoblastoma cells compared with hTERT-RPE retinal pigment epithelial cells. shRNA-mediated stable silencing of UNC5D markedly suppressed cell proliferation, colony formation, migration, and invasion, while promoting apoptosis in both Rb cell lines. In a nude mouse xenograft model, UNC5D knockdown significantly reduced tumor growth, confirming its pro-tumorigenic role in vivo. To elucidate the underlying mechanism, whole-transcriptome sequencing was performed following UNC5D depletion, revealing significant enrichment of downregulated genes associated with PI3K/AKT signaling. Consistently, phosphorylation of PI3K and AKT was markedly reduced in UNC5D-silenced cells and xenograft tumors, whereas total PI3K and AKT protein levels remained largely unchanged. Because UNC5 family receptors are known to interact with netrin ligands, we further investigated the relationship between UNC5D and netrin-1 (NTN1). Correlation analysis, molecular docking, co-immunoprecipitation, and expression studies demonstrated a strong association between UNC5D and NTN1 in retinoblastoma cells. Collectively, our findings identify UNC5D as a previously unrecognized oncogenic regulator of retinoblastoma and suggest that UNC5D is associated with NTN1 signaling and contributes to PI3K/AKT pathway activation and Rb tumor progression. These results suggest the UNC5D-NTN1-PI3K/AKT axis as a potential therapeutic target for retinoblastoma.

## 1. Introduction

Retinoblastoma (Rb) is the most common intraocular malignancy of childhood and accounts for approximately 2–4% of all pediatric cancers worldwide [1]. Although biallelic inactivation of the RB1 tumor suppressor gene represents the initiating event in most cases, accumulating evidence indicates that additional genetic and epigenetic alterations are required for tumor progression and malignant transformation [2]. Despite significant advances in focal therapies, intra-arterial and intravitreal chemotherapy, and systemic treatment regimens, vision loss, recurrence, and extraocular dissemination remain important clinical challenges [3,4]. A better understanding of the molecular mechanisms underlying Rb progression is therefore essential for the development of targeted therapeutic strategies.

Recent studies have highlighted the contribution of aberrant signaling pathways and epigenetic dysregulation to Rb pathogenesis [5]. Through an integrative analysis of DNA methylation, miRNA, and mRNA expression datasets, we previously identified UNC5D as a candidate regulator consistently dysregulated in retinoblastoma [6]. UNC5D (Uncoordinated-5 Netrin Receptor D) belongs to the UNC5 family of dependence receptors and functions as a transmembrane receptor for netrin ligands [7]. Structurally, UNC5D contains extracellular immunoglobulin-like and thrombospondin domains, together with intracellular ZU5 and death domains that mediate downstream signaling [7]. Beyond its established roles in axonal guidance, neuronal migration, and tissue development, UNC5D has been implicated in the regulation of cell survival and apoptosis [8–11]. In the absence of its ligand, UNC5D functions as a dependence receptor capable of triggering apoptotic signaling, whereas ligand engagement promotes cell survival and migration [11].

The role of UNC5D in cancer remains incompletely understood and appears highly context dependent. Previous studies have reported tumor-suppressive functions of UNC5D in several malignancies, including renal cell carcinoma, prostate cancer, thyroid cancer, breast cancer, and hepatocellular carcinoma, where its expression is frequently reduced through promoter hypermethylation or loss of heterozygosity [12–16]. Conversely, our earlier study demonstrated significant upregulation of UNC5D at both the mRNA and protein levels in Y79 and Weri-Rb1 retinoblastoma cells compared with hTERT-RPE cells [6], suggesting a distinct and previously unrecognized role in Rb biology.

In the present study, we investigated the functional significance of UNC5D in retinoblastoma using in vitro and in vivo models. Stable shRNA-mediated silencing of UNC5D was employed to examine its effects on proliferation, apoptosis, migration, invasion, and tumorigenicity. Transcriptomic profiling combined with molecular and biochemical analyses identified PI3K/AKT signaling as a major pathway regulated by UNC5D. Furthermore, interaction studies revealed a close association between UNC5D and its ligand netrin-1 (NTN1) in retinoblastoma cells. Collectively, our findings identify UNC5D as a novel oncogenic regulator in retinoblastoma and provide evidence that UNC5D is extensively associated with NTN1 signaling and contributes to PI3K/AKT pathway activation and Rb tumor progression.

## 2. Materials and Methods

### 2.1 Cell culture

Human retinoblastoma cell lines Y79 (RCB1645) and WERI-Rb-1 (HTB-169), and the hTERT-immortalized retinal pigment epithelial cell line hTERT-RPE-1 (CRL-4000), were obtained from RIKEN BioResource Center and the American Type Culture Collection (ATCC), respectively. Y79 and WERI-Rb-1 cells were cultured in RPMI-1640 medium (Gibco, Thermo Fisher Scientific) supplemented with 10% fetal bovine serum (FBS), L-glutamine, and antibiotic-antimycotic solution. hTERT-RPE-1 cells were maintained in DMEM/F12 medium supplemented with 10% FBS and 0.01 mg/ml hygromycin B (Sigma-Aldrich). Cells were cultured at 37°C in a humidified atmosphere containing 5% CO₂. All cell lines were routinely tested and confirmed to be free of mycoplasma contamination.

### 2.2 Generation of stable UNC5D-knockdown cell lines

Stable UNC5D knockdown was achieved using shRNA constructs targeting human UNC5D (VectorBuilder, USA). A scrambled shRNA construct served as a negative control. Cells were transfected using Lipofectamine 2000 (Invitrogen) according to the manufacturer’s instructions and selected with puromycin (2 μg/ml) to generate stable cell lines. Knockdown efficiency was confirmed by quantitative real-time PCR (qRT-PCR) and western blotting. shRNA sequences and vector maps are provided in **Supplementary Table S1** and **Supplementary Fig. S1**.

### 2.3 Quantitative real-time PCR

Total RNA was isolated using RNAiso Plus (TaKaRa) and reverse-transcribed using the PrimeScript First Strand cDNA Synthesis Kit (TaKaRa). qRT-PCR was performed using TB Green Premix Ex Taq II (TaKaRa). Relative gene expression was calculated using the 2^−ΔΔCt method with GAPDH as the internal control. Primer sequences are listed in **Supplementary Table S2**.

### 2.4 Western blot analysis

Total protein was extracted using RIPA lysis buffer supplemented with protease inhibitors. Protein concentrations were determined using a BCA Protein Assay Kit (Thermo Scientific). Equal amounts of protein were resolved by SDS-PAGE, transferred to nitrocellulose membranes, and incubated with primary antibodies overnight at 4°C followed by HRP-conjugated secondary antibodies. Signals were detected using enhanced chemiluminescence (Bio-Rad) and quantified using ImageJ (Fiji v1.54f). Antibodies used in this study are listed in **Supplementary Table S3**.

### 2.5 Cell proliferation

Following transfection and generation of stable cell lines, approximately 3×10^3^ cells were seeded in each well of a 96-well plate and cell proliferation was assessed using the Cell Counting Kit-8 (CCK-8) (96992, Sigma) according to the manufacturer’s protocol. For cell proliferation assay, at 0, 24, 48 and 72 hours post culture, the cells were incubated with 10μl of the CCK-8 solution per well for 4 hours in the 37℃ incubator. The absorbance was then taken at 450 nm. in Multiskan SkyHigh microplate reader (Thermo Scientific, USA).

### 2.6 Colony formation assays

Anchorage-independent growth was evaluated using a soft agar colony formation assay. A total volume of 1 ml. of 0.7% agar in the complete medium was evenly spread in each well of a 6-well plate. Following transfection, approximately 7000 cells were seeded into the top layer of the base agar after being suspended in 1 ml. of 0.3% agar in complete medium. The cells were fed with 500 µl of complete medium twice weekly for four weeks. After being fixed with 70% ethanol for 10 min at room temperature, the colonies in soft agar were stained with 0.05% methylene blue for 20 min. The pictures were then captured using a light microscope. Colony numbers were quantified using ImageJ Fiji (1.54f) software.

### 2.7 Transwell migration and invasion assays

For migration assay, approximately 3×10^4^ transfected Y79 & Weri Rb1 cells were suspended in 200μl serum-free RPMI media and put into transwell inserts (pore size-8μm) (3464, Corning). All transwell inserts were placed in 24-well plates with 500μl RPMI supplemented with 20% FBS underneath each well. Then these plates were incubated in the 37°C incubator for 12-15 h. Cells that failed to migrate across the semi-permeable membrane of the transwell inserts were then carefully scraped off with a cotton bud. The migrated cells on the lower portion of the inserts were fixed with methanol and acetic acid (3:1) mixture for 40 min at room temperature. After fixation, the cells were stained with 0.5% methylene blue for 30 min and counted under a light microscope. Similar procedures were followed for the invasion assay but the matrigel-coated transwell inserts (354480, Corning) were used.

### 2.8 Cell cycle and apoptosis analyses

Transfected Y79 and Weri Rb1 cells were harvested and seeded in a 6 well ultra-low attachment plate (3471, Corning). Approximately, 1×10^6^ no. of cells were collected and washed twice with ice-cold PBS. After that, the cells were fixed overnight with 70% ethanol at 4°C. Following fixation, the cells underwent two more rounds of ice-cold PBS washing before being treated with 1 mg/ml RNaseA (19101, Qiagen) for 30 minutes at 37°C. Then the cells were stained with 50μg/ml propidium iodide (P4170, Sigma) in the dark for 30 min. Flow cytometric analysis was done using a BD LSR Fortessa (BD Biosciences, USA) and the results were analyzed by using FlowJo v10.8.1 (FlowJo LLC, USA).

The cellular apoptosis rate was determined using an Annexin V-APC Apoptosis detection kit (550475, BD Biosciences, USA). Transfected Y79 and Weri Rb1 cells were cultured and approximately 1×10^6^ no. of cells were washed with 1X PBS. Then the cells were resuspended in 500μl of 1X Annexin-V-binding buffer (BD Pharmingen, USA) and 5μl of each of Annexin V-APC and propidium iodide (PI) staining solution were added. The samples were then incubated for 15 min at room temperature in the dark. The stained cells were subsequently evaluated for the detection of apoptosis using a BD LSR Fortessa (BD Biosciences, USA). Cells classified as early apoptotic (Q3 quadrant) were those that tested positive for Annexin V-APC but negative for PI staining. Data were analyzed using FlowJo v10.8.1 software (FlowJo LLC, USA).

### 2.9 RNA sequencing and bioinformatic analysis

Total RNA was extracted from wild-type, shNC, and shUNC5D Y79 cells using the RNeasy Mini Kit (Qiagen). Libraries were prepared using the NEBNext Ultra II RNA Library Preparation Kit and sequenced on an IlluminaNovaSeq 6000 platform (2 × 150 bp). Clean reads were aligned to the human reference genome (GRCh38) using STAR, and gene counts were generated using featureCounts. Differential expression analysis was performed using DESeq2. Genes with p < 0.05 and |log₂ fold change| ≥ 2 were considered differentially expressed. Gene Ontology, KEGG pathway enrichment, and gene set enrichment analyses (GSEA) were performed using ShinyGO and clusterProfiler. RNA sequencing data have been deposited in the Gene Expression Omnibus under accession number GSE306604. Detailed protocol for RNA sequencing is provided in **Supplementary method 3**.

### 2.10 Xenograft tumor model

All animal experiments were approved by the Institutional Animal Ethics Committee of CSIR–Indian Institute of Chemical Biology (Approval No. IICB/AEC/JAN/2024/12) and conducted in accordance with institutional guidelines. The care of the animals was provided following the guidelines of CSIR-Indian Institute of Chemical Biology, Kolkata. Six to eight weeks old female athymic CrTac:NCr-Foxn^1nu^ nude mice (weight range: 20-25 gm) were obtained from ACTREC, Mumbai. The institutional animal facility provided housing and husbandry conditions. A maximum of five mice were kept in each Individually Ventilated Cage (IVC) with corncob bedding. Every mouse was housed in a 12-hour light/dark cycle with unrestricted access to foodstuff and drink. All mice were assigned to the shNC and shUNC5D groups (n = 5 per group) prior to injection. Then they were subcutaneously injected with 5 × 10⁶ Y79 cells expressing either control shRNA or UNC5D-targeting shRNA suspended in matrigel. All mice were tracked to monitor the overall health status including body weight and tumor volume over the course of the entire study. Randomization and blinding have been applied to create groups of equal size. Tumor growth was monitored every four days, and tumor volume was calculated using the formula: volume = (length × width²)/2. Tumor growth curves were analyzed using two-way ANOVA, whereas final tumor weight were compared using unpaired Student’s t-tests. Mice were euthanized on day 15, and tumors were collected for histological and molecular analyses.

### 2.11 Immunohistochemistry

Tumor tissues were fixed in 4% paraformaldehyde, cryoprotected, embedded in OCT compound, and sectioned at 10 μm thickness. Immunohistochemistry was performed using an HRP/DAB detection kit (ab64261, Abcam) with antibodies against UNC5D and Ki67. The anti-UNC5D primary antibody (Bioss) was used at a 1:500 dilution and anti-Ki67 primary antibody (CST) was used at a 1:600 dilution. A biotinylated secondary antibody provided with the above mentioned IHC kit was used at a 1:400 dilution. Tissue sections on the slides were then stained with DAB (3,3′-diaminobenzidine) and counterstained with hematoxylin (Himedia, India). Images were acquired using an Olympus microscope and quantified using ImageJ Fiji (1.54f) software.

### 2.12 Immunofluorescence

For immunofluorescence, cells were fixed, permeabilized, and incubated with anti-NTN1 primary antibody (A16236, ABclonal) followed by Alexa Fluor-conjugated secondary antibodies (Invitrogen, ThermoFisher Scientific). Nuclei were counterstained with Hoechst 33342 (H3570, Invitrogen, ThermoFisher Scientific). Images were captured using a Zeiss LSM980 confocal microscope and analyzed using ZEN software.

### 2.13 Correlation analyses

The correlation between UNC5D and NTN1 in normal human retina was analyzed by using the Correlation AnalyzeR software (https://github.com/Bishop-Laboratory/correlationAnalyzeR). This software is mainly used for exploration of co-expression correlations, prediction of gene–gene relationships, gene functions, and gene set topology. The Pearson correlation coefficient (R) value of +1 indicates a perfect positive co-relationship whereas −1 indicates a perfect negative co-relationship between two genes.

### 2.14 Molecular docking

To check the binding efficiency between UNC5D and NTN1, molecular docking was performed by using the HDOCK software (http://hdock.phys.hust.edu.cn/). In this study, the model for UNC5D has been taken from RCSB PDB (ID- 5FTU)- (Tetrameric complex of Latrophilin 3, UNC5D & FLRT2). The model for NTN1 was taken from RCSB PDB (ID- 6FKQ)- (The crystal structure of a fragment of netrin1 in complex with a fragment of Draxin).

### 2.15 Construction of PPI network

Search Tools For Retrieval of Interacting Genes/ Proteins (STRING) (https://string-db.org/) database was employed to investigate the potential protein-protein interaction between UNC5D and NTN1.

### 2.16 Co-immunoprecipitation

For co-immunoprecipitation assay, approximately 500 μg of protein sample was kept in lysis buffer (50 mM Tris HCl pH 7.4, 150 mM NaCl, 1% NP-40, 5% glycerol, 1 mM EDTA, 0.1% Tween-20) and washed with equilibrated Protein A agarose beads (20333, Pierce, Thermo Scientific). After that, ∼25μl beads were pre-incubated for 3-4 hours at 4°C with the corresponding primary antibodies (1:100). An equivalent volume of cleared protein lysate was subsequently added to the beads-IP antibody mixture and beads-IgG mixture, incubated overnight under rotary agitation at 4℃. Next day, the samples were washed repeatedly with wash buffer (1X PBS supplemented with 0.05% Triton-X-100). Following a 5-minute centrifugation at 2000g at 4°C, the supernatant was collected, boiled with 5X Laemmli lysis buffer, and subjected to western blot assay. For western blot assay, 20 μL of the eluted immunoprecipitate or 50 μg of cell lysate was separated by SDS-PAGE. The details of the antibodies used were listed in **Supplementary Table S3**.

### 2.17 Statistical analysis

All experiments were performed using at least three independent biological replicates unless otherwise stated. Data are presented as mean ± standard deviation (SD). The number of biological replicates (n) is specified in each figure legend. SD error bars are used to represent the estimates of variation within each group. Statistical analyses were performed using GraphPad Prism v8.0.2 (GraphPad Software, USA) and R software (v4.5.1) (http://www.r-project.org).

Comparisons between two groups were performed using two-tailed unpaired Student’s t-tests. For xenograft tumor growth studies, statistical significance was determined using two-way analysis of variance (ANOVA) followed by Sidak’s multiple-comparison test. Cell proliferation assays, cell cycle distribution, apoptosis assays, qRT-PCR, western blot densitometry, colony formation, migration, invasion, and immunohistochemical quantification were analyzed using unpaired Student’s t-tests. A p value < 0.05 was considered statistically significant.

Statistical significance is indicated as follows: p < 0.05 (*), p < 0.01 (**), p < 0.001 (***), and p < 0.0001 (****).

## 3. Results

### 3.1 UNC5D is upregulated in retinoblastoma and promotes malignant phenotypes in vitro

Analysis of publicly available TCGA datasets using GEPIA and UALCAN revealed elevated UNC5D expression across eight of 33 cancer types, including lower-grade glioma and glioblastoma multiforme (Fig. S2A, B). Consistent with these observations, UNC5D protein expression was higher in HeLa, SH-SY5Y, and HuH7 cancer cell lines than in non-transformed HEK293T cells (Fig. S2C). Our previous integrative multi-omics analysis similarly identified UNC5D as significantly upregulated in retinoblastoma (Rb) cells relative to hTERT-RPE cells [6]. Because retinoblastoma is believed to originate from cone precursor-like cells [20,23,24], we further analyzed single-cell RNA-sequencing data (GSE249995), which demonstrated preferential UNC5D expression within cone precursor-like cell clusters (Fig. S3A).

To investigate the functional role of UNC5D in Rb, stable UNC5D-knockdown Y79 and Weri-Rb1 cell lines were established using shRNA. Efficient silencing was confirmed by qRT-PCR and immunoblotting (Fig. 1A, B). UNC5D depletion significantly reduced cell proliferation from 48 h onward in both cell lines (Fig. 1C). Consistent with these findings, soft agar assays demonstrated a marked reduction in clonogenic growth following UNC5D knockdown (Fig. 1D). Moreover, Transwell assays showed significantly impaired migratory and invasive capacities in UNC5D-silenced Y79 and Weri-Rb1 cells compared with scramble controls (Fig. 1E, F). Together, these findings indicate that UNC5D promotes proliferative, clonogenic, migratory, and invasive phenotypes in retinoblastoma cells.

**Figure 1.**
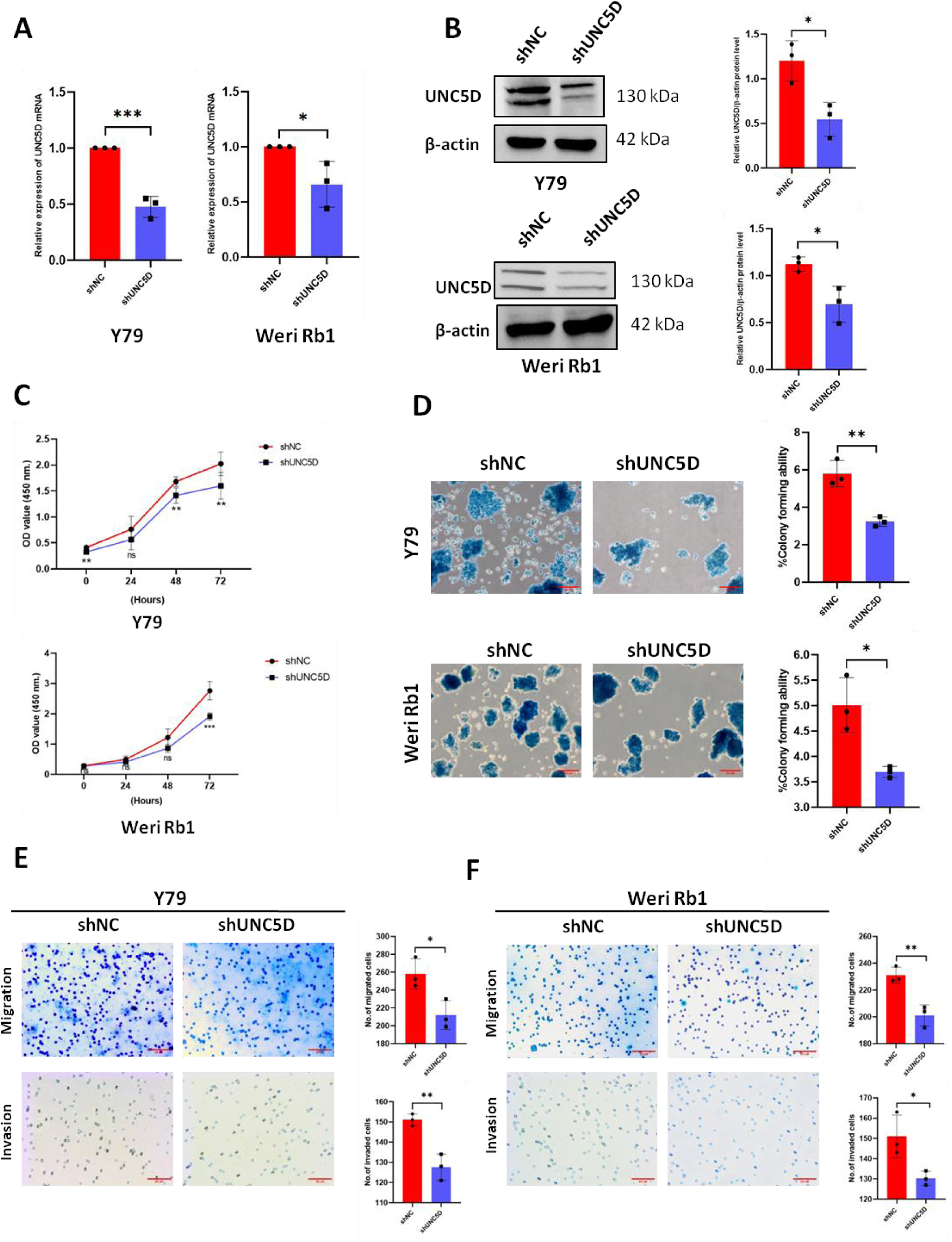
UNC5D regulates the oncogenic properties of retinoblastoma cells *in vitro.* (A) qRT-PCR & (B) western blot assays confirmed the knockdown of UNC5D in Y79 & Weri Rb1 cells as compared to the cells treated with scramble control (n=3). Respective densitometry plots of the proein blots [*p˂0.05; **p˂0.01 and ***p˂0.001]. (C) Investigating the proliferation of Y79 & Weri Rb1 retinoblastoma cells using CCK8 assay at different time intervals after shRNA mediated knockdown of UNC5D. (D) Soft agar colony formation assay to investigate the colony forming ability of Y79 & Weri Rb1 cells. Images were taken in a light microscope with 40X magnification (Scale bars, 20 μm) (E) Transwell migration and invasion assays in Y79 cells. (F) Transwell migration and invasion assays in Weri Rb1 cells. Images were taken in a light microscope with 10X magnification (Scale bars, 20 μm). All data are represented as mean ± SD (n=3) [*p˂0.05; **p˂0.01 and ***p˂0.001].

### 3.2 UNC5D facilitates cell-cycle progression and suppresses apoptosis in retinoblastoma cells

To determine whether the reduced proliferative capacity observed following UNC5D depletion was associated with altered cell-cycle progression, flow cytometric analysis was performed. UNC5D knockdown significantly decreased the proportion of cells in S phase in both Y79 and Weri-Rb1 cells, accompanied by a modest increase in the G1 population (Fig. 2A), indicating impaired cell-cycle progression. Consistent with these observations, Cyclin D1 protein expression was markedly reduced following UNC5D silencing (Fig. 2C, D).

**Figure 2.**
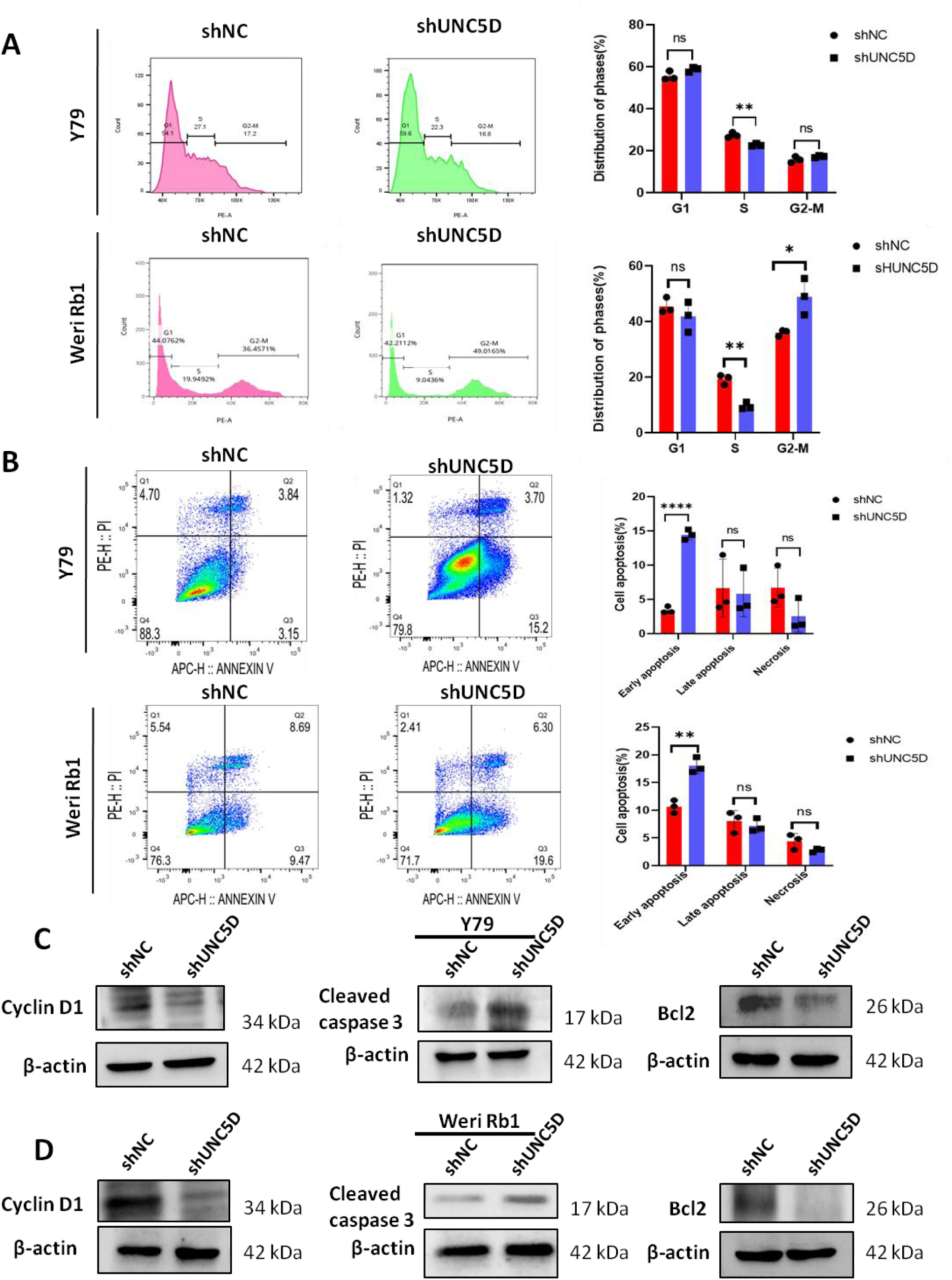
UNC5D promotes cell cycle progression and deactivates apoptotic machinery in retinoblastoma. (A) Analysis of rate of different stages of cell cycle in scramble control (shNC) and UNC5D knocked down (shUNC5D) Y79 & Weri Rb1 cells. (B) Analysis of rate of apoptosis in scramble control and UNC5D knocked down Y79 & Weri Rb1 cells by Annexin-V-APC-PI assay. (C) Investigating the relative expressions of the cell cycle related protein cyclin D1 and apoptosis related proteins cleaved caspase 3 & Bcl2 in scramble control and UNC5D knocked down Y79 cells. (D) Investigating the relative expressions of the cell cycle related protein cyclin D1 and apoptosis related proteins cleaved caspase 3 & Bcl2 in scramble control and UNC5D knocked down Weri Rb1 cells. All data are represented as mean ± SD (n=3) [*p˂0.05; **p˂0.01 and ***p˂0.001].

Annexin V-APC/PI staining further demonstrated that depletion of UNC5D significantly increased the proportion of early apoptotic cells in both Rb cell lines compared with scramble controls (Fig. 2B). This phenotype was accompanied by increased expression of cleaved caspase-3 and reduced expression of the anti-apoptotic protein Bcl-2 (Fig. 2C, D), confirming activation of apoptotic signaling. Collectively, these results indicate that UNC5D supports retinoblastoma cell survival by promoting cell-cycle progression while suppressing apoptosis.

### 3.3 UNC5D activates PI3K/AKT signaling in retinoblastoma

To elucidate the molecular mechanisms underlying UNC5D-mediated retinoblastoma progression, we performed whole-transcriptome sequencing of Y79 cells following UNC5D knockdown. Differential expression analysis identified 173 significantly upregulated and 423 significantly downregulated genes (P < 0.05, |log₂FC| ≥ 2) (Fig. S4C). KEGG pathway enrichment analysis demonstrated that upregulated genes were primarily associated with extracellular matrix (ECM)-receptor interaction and focal adhesion pathways (Fig. 3A), whereas downregulated genes were significantly enriched in calcium signaling, MAPK signaling, and PI3K/AKT signaling pathways (Fig. 3B).

**Figure 3.**
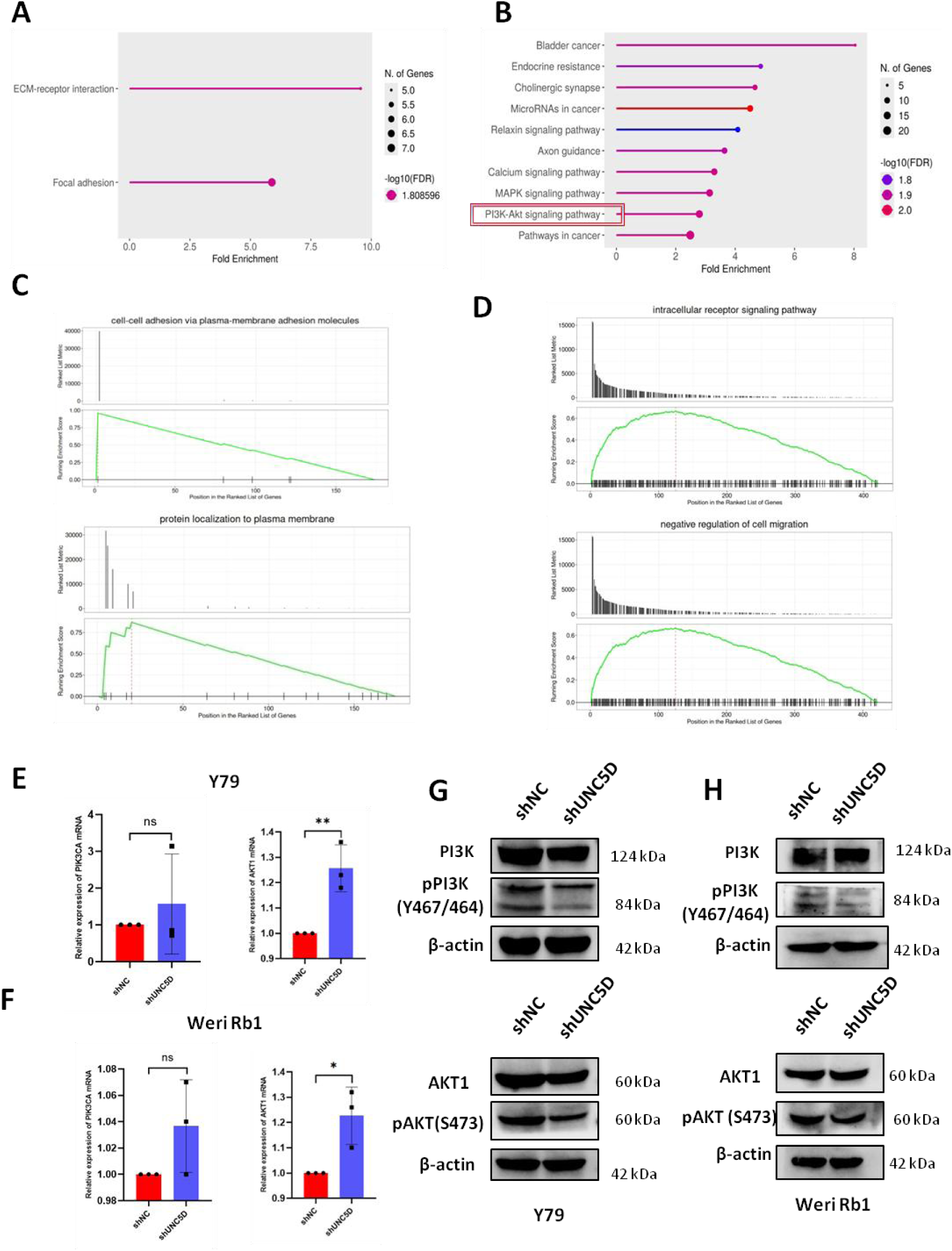
Alteration in PI3K/AKT signaling pathway in response to UNC5D downregulation in retinoblastoma.(A) KEGG Pathway enrichment analysis of upregulated; [p<0.05, log_2_FC>+-2] genes (in shNC & shUNC5D groups). (B) KEGG Pathway enrichment analysis of downregulated; [p<0.05, log_2_FC>+-2] genes (in shNC & shUNC5D groups). (C) Representative GSEA showing that upregulated genes were enriched in cell-cell adhesion via plasma membrane adhesion molecules (NES=1.38, p=0.01, q=0.98) and protein localization to plasma membrane (NES=1.35, p=0.03, q=0.98) in shNC & shUNC5D groups. (D) Representative GSEA showing that downregulated genes were enriched in intracellular receptor signaling pathway (NES=1.51, p=0.009, q=0.69) and negative regulation of cell migration (NES=1.52, p=0.02, q=0.69) in shNC & shUNC5D groups. (E) Investigating the relative expressions of PI3KCA & AKT1 mRNAs in shNC & shUNC5D groups of Y79 cells by qRT-PCR. (F) Investigating the relative expressions of PI3KCA & AKT1 mRNAs in shNC & shUNC5D groups of Weri Rb1 cells by qRT-PCR. (G) Investigating the relative expressions of the cell proliferation marker proteins viz. PI3K, p-PI3K, AKT1 & p-AKT in shNC & shUNC5D groups of Y79 cells by western blotting. (H) Investigating the relative expressions of the cell proliferation marker proteins viz. PI3K, p-PI3K, AKT1 & p-AKT in shNC & shUNC5D groups of Weri Rb1 cells by western blotting. All data are represented as mean ± SD (n=3) [*p˂0.05; **p˂0.01 and ***p˂0.001].

Gene set enrichment analysis (GSEA) further identified nominal enrichment of biological processes related to cell-cell adhesion via plasma membrane adhesion molecules and protein localization to the plasma membrane among upregulated genes, whereas intracellular receptor signaling pathway and negative regulation of cell migration were enriched among downregulated genes following UNC5D silencing (Fig. 3C, D). Gene Ontology analyses of biological processes, cellular components, and molecular functions, together with volcano plots and heat maps of differentially expressed genes, are presented in Figs. S4–S6.

Given the prominent enrichment of the PI3K/AKT pathway, we next validated key signaling components experimentally. Quantitative RT-PCR showed no significant change in PIK3CA mRNA expression following UNC5D knockdown, whereas AKT1 mRNA expression was modestly increased in the shUNC5D group as compared to the shNC groups in both Y79 and Weri-Rb1 cells (Fig. 3E, F). In contrast, immunoblot analysis demonstrated no significant differences in total PI3K or AKT protein abundance but revealed a marked reduction in phosphorylated PI3K (p-PI3K) and phosphorylated AKT (p-AKT) in UNC5D-silenced cells (Fig. 3G, H). These findings indicate that UNC5D regulates retinoblastoma progression primarily by promoting activation of the PI3K/AKT signaling pathway rather than altering total PI3K or AKT expression.

### 3.4 UNC5D promotes retinoblastoma tumor growth in vivo

To determine whether UNC5D contributes to retinoblastoma progression in vivo, Y79 cells expressing control (shNC) or UNC5D-targeting shRNA (shUNC5D) were implanted subcutaneously into athymic nude mice. Tumors derived from UNC5D-silenced cells exhibited significantly reduced growth compared with controls, resulting in markedly lower tumor weight and tumor volume at the experimental endpoint (Fig. 4B–D).

**Figure 4.**
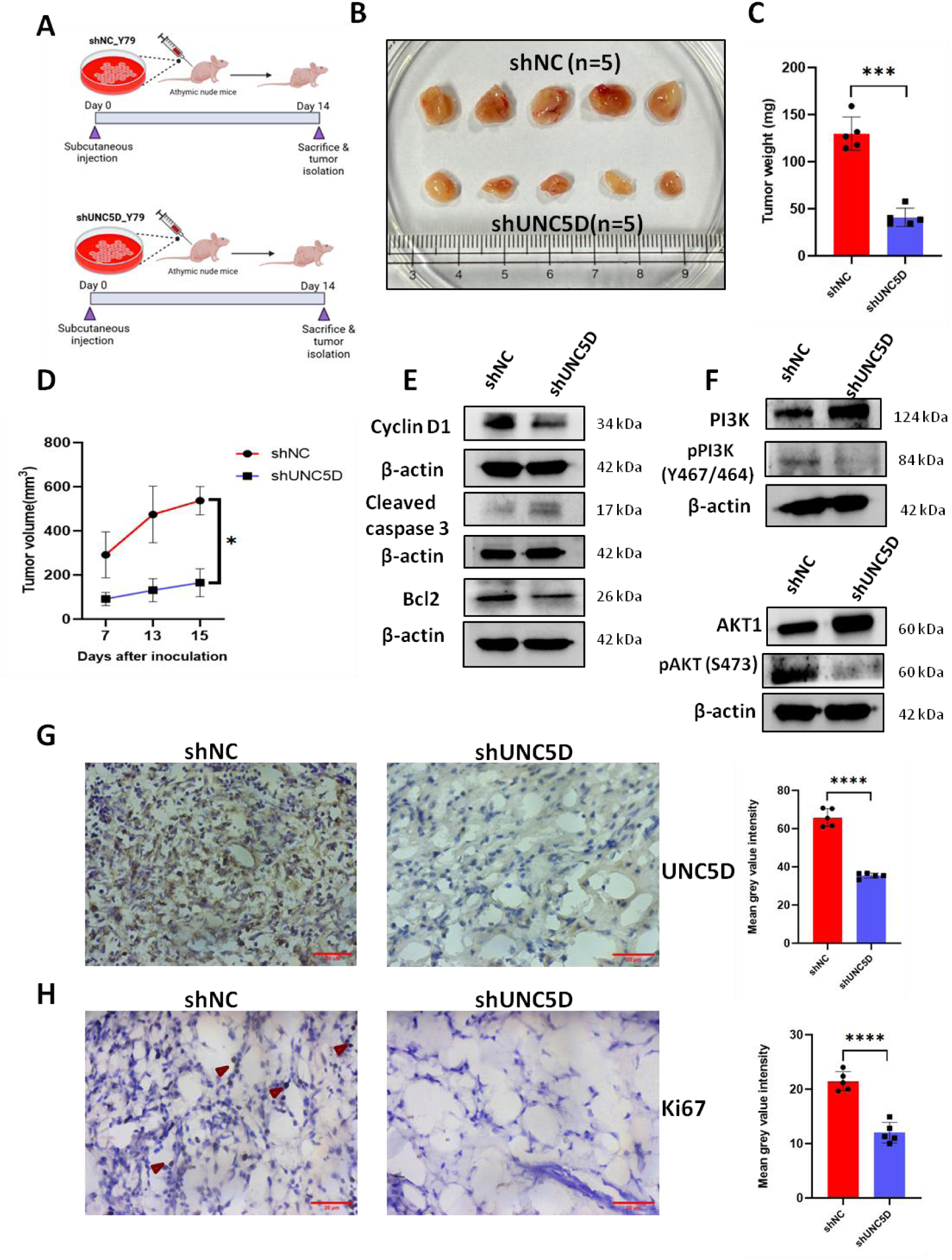
Silencing of UNC5D represses tumorigenesis of retinoblastoma cells *in vivo.* (A) Schematic illustrating the *in vivo* experimental design. (Made with Biorender.com). B. Representative images of isolated subcutaneous tumors in mice receiving Y79 scramble control cells and cells treated with shUNC5D. Quantification of tumor weight (C) and volume (D). Data are shown as the mean ± SD (n = 5). [*p* < 0.05(*), *p* < 0.01(**), p> 0.001(***) and p> 0.0001(****)]. (E) Investigating the relative expressions of the cell cycle related protein cyclin D1 and apoptosis related proteins cleaved caspase 3 & Bcl2 in protein lysates isolated from xenograft tumors (both shNC & shUNC5D groups). (F) Investigating the relative expressions of the cell proliferation marker proteins PI3K, p-PI3K, AKT1 & p-AKT in protein lysates isolated from xenograft tumors (both shNC & shUNC5D groups). (G) UNC5D knockdown confirmed by Immunohistochemical staining (shNC & shUNC5D groups) (Magnification, 40×) (Scale bars, 20 μm). (H) Ki67 staining of tumor tissues (shNC & shUNC5D groups) (Magnification, 40×) (Scale bars, 20 μm). Ki67 positive cells were indicated by arrowheads in the shNC tissue panel. Data are shown as the mean ± SD (n = 5). [*p* < 0.05(*), *p* < 0.01(**), p> 0.001(***)].

Immunoblot analysis of xenograft tumor lysates demonstrated reduced expression of Cyclin D1, Bcl-2, phospho-PI3K (p-PI3K), and phospho-AKT (p-AKT), together with increased cleaved caspase-3 expression following UNC5D depletion (Fig. 4E, F). Consistent with these findings, immunohistochemical analysis confirmed reduced UNC5D and Ki67 expression in shUNC5D tumors compared with shNC controls (Fig. 4G, H). Representative hematoxylin and eosin (H&E)-stained sections are shown in Fig. S7A, B. Collectively, these findings demonstrate that UNC5D promotes retinoblastoma tumor growth in vivo by enhancing cell proliferation, activating PI3K/AKT signaling, and suppressing apoptosis.

### 3.5 UNC5D associates with NTN1 in retinoblastoma cells

Because UNC5 family receptors function through interactions with netrin ligands [9], we next investigated the relationship between UNC5D and NTN1 in retinoblastoma. Correlation analysis of normal human retinal tissue demonstrated a strong positive association between UNC5D and NTN1 expression (R = 0.77) (Fig. 5A). Protein-protein docking further predicted a favorable interaction between UNC5D and NTN1 (HDOCK docking score = −294; confidence score = 0.95) (Fig. 5B), while the structural models used for docking are provided in Fig. S8A, B.

**Figure 5.**
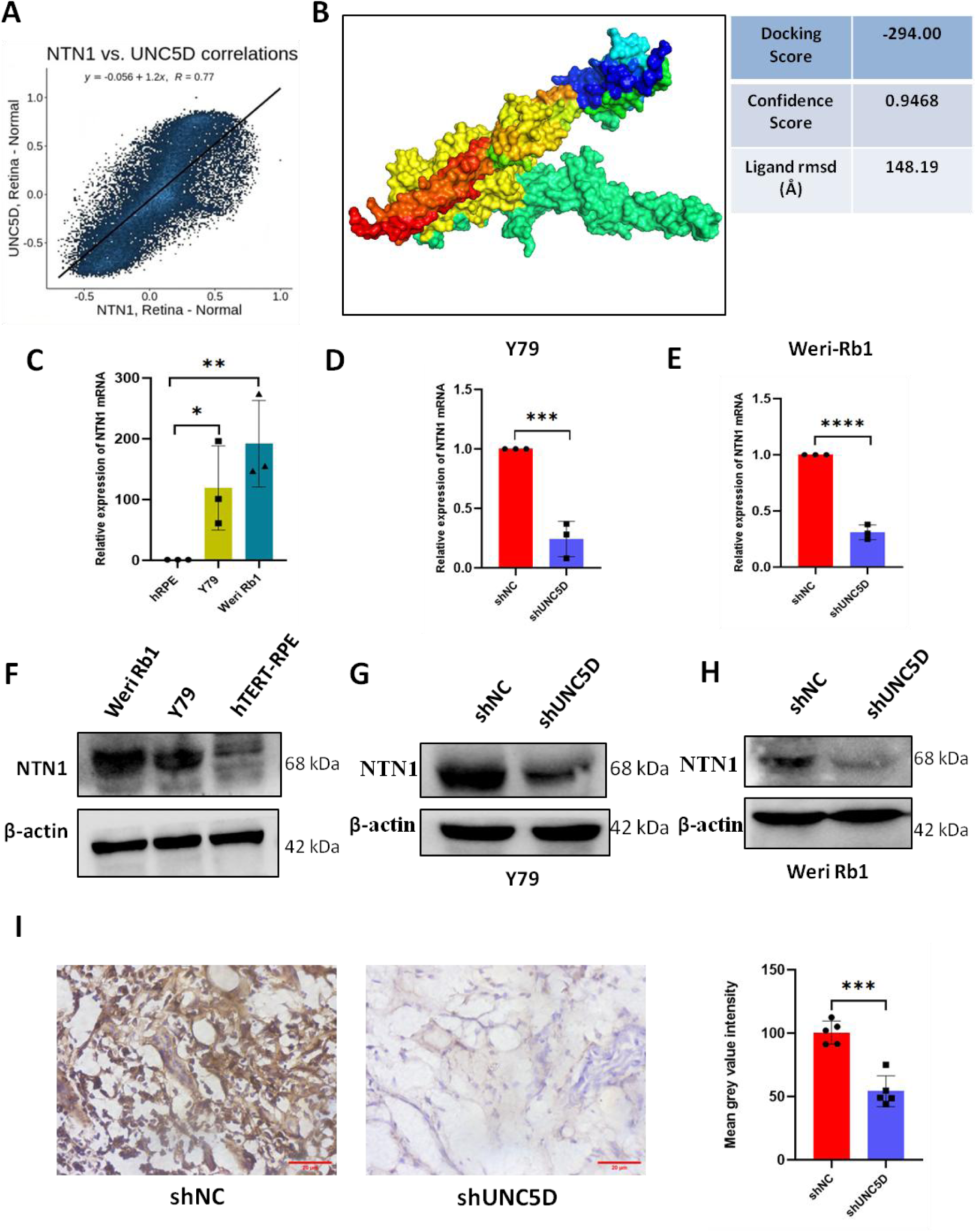
NTN1 and UNC5D interact strongly to promote Rb progression. (A) Correlation between NTN1 & UNC5D in normal human retina. (B) Protein-protein docking model and docking score of UNC5D & NTN1. (C) Investigating the relative expression of NTN1 mRNA in hTERT-RPE cells and Y79 & Weri Rb1 Rb cells by qRT-PCR (n=3) [*p˂0.05; **p˂0.01 and ***p˂0.001]. (D) Investigating the relative expression of NTN1 mRNA in shNC & shUNC5D Y79 cells by qRT-PCR (n=3) [*p˂0.05; **p˂0.01 and ***p˂0.001]. (E) Investigating the relative expression of NTN1 mRNA in shNC & shUNC5D Weri Rb1 cells by qRT-PCR (n=3) [*p˂0.05; **p˂0.01 and ***p˂0.001]. (F) Investigating the relative expression of NTN1 protein in Y79 & Weri Rb1 Rb cells and hTERT-RPE cells by western blotting. (G) Investigating the relative expression of NTN1 protein in negative control & UNC5D knockdown Y79 cells by western blotting. (H) Investigating the relative expression of NTN1 protein in negative control & UNC5D knockdown Weri Rb1 cells by western blotting. (I) Immunohistochemical staining of NTN1 in negative control and UNC5D-knockdown groups of xenograft tumor tissues (Magnification, 40×) (Scale bars, 20 μm). Data are shown as the mean ± SD (n = 5). [*p* < 0.05(*), *p* < 0.01(**), p> 0.001(***)].

NTN1 expression was subsequently evaluated in retinoblastoma cells. Both NTN1 mRNA and protein levels were elevated in Y79 and Weri-Rb1 cells relative to hTERT-RPE cells (Fig. 5C, F). Silencing UNC5D significantly reduced NTN1 mRNA (Fig. 5D, E) and protein expressions (Fig. 5G, H) in both retinoblastoma cell lines and decreased protein expression in xenograft tumor tissues (Fig. 5I). Consistent with these findings, immunofluorescence analysis demonstrated decreased NTN1 expression in UNC5D-depleted Y79 cells (Fig. 6A).

**Figure 6.**
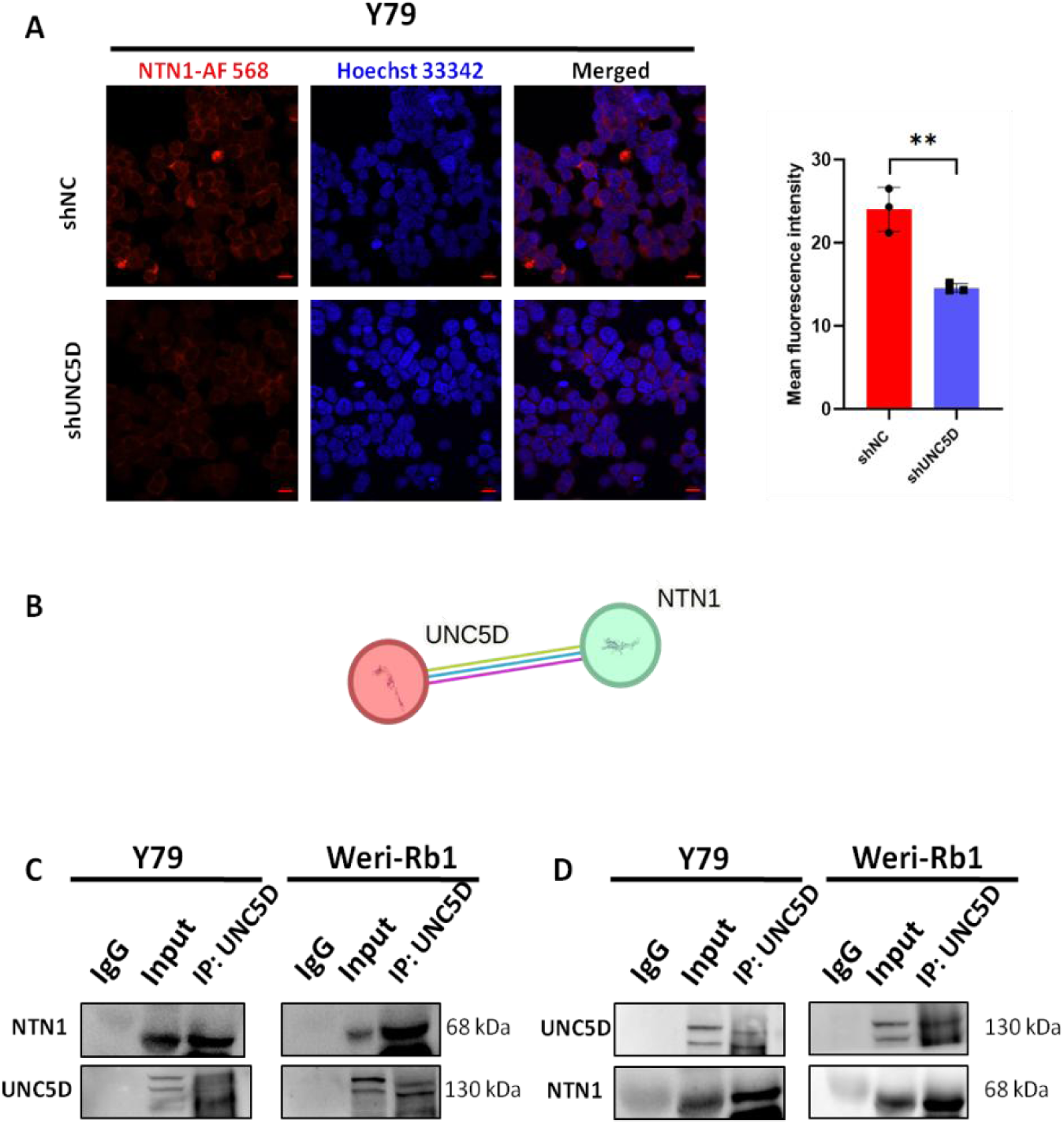
NTN1 strongly interacts with UNC5D to induce Rb pathogenesis. (A) Confocal microscopy images showing shNC group and shUNC5D group of Y79 cells, stained for NTN1 (Alexa fluor 568, red) and nucleus (DAPI, blue) after silencing UNC5D. shNC group of cells were used as control. All the images were quantified using Zeiss ZEN lite software (Scale bars, 10 μm). All data are represented as mean ± SD (n=3) [*p˂0.05; **p˂0.01 and ***p˂0.001].(B) PPI network showing the interaction between UNC5D & NTN1 proteins using STRING database. (C) Immunoprecipitation of UNC5D and the blots were probed for NTN1 in Y79 and Weri Rb1 Rb cells. (D) Immunoprecipitation of NTN1 and the blots were probed for UNC5D in Y79 and Weri Rb1 Rb cells. Whole cell lysate (input) was used as positive control while IgG was considered as negative control.

**Figure 7.**
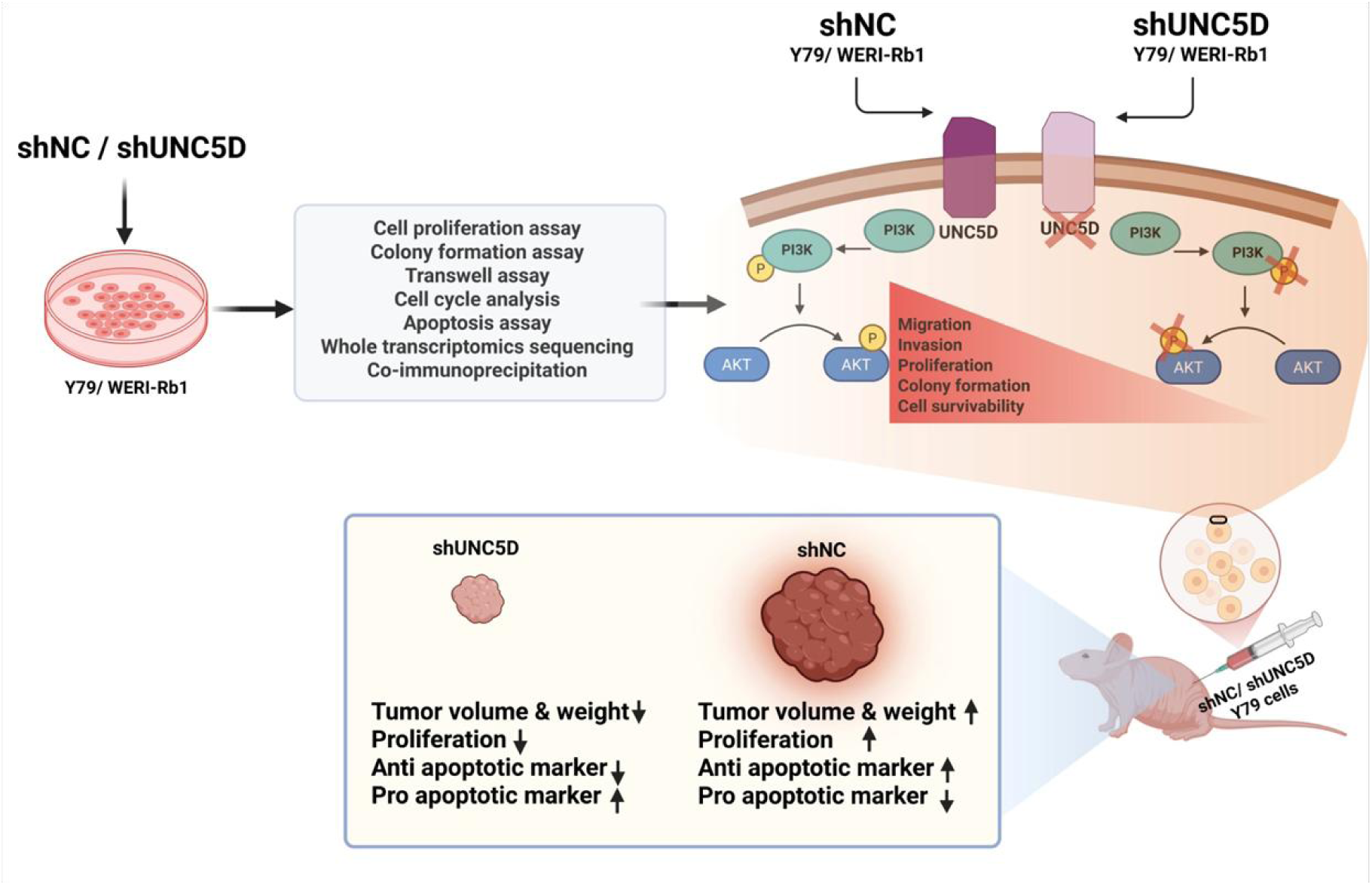
A schematic representation of the overall study. UNC5D plays a key oncogenic role by promoting retinoblastoma progression via activating the PI3K/AKT signaling pathway. Knockdown of UNC5D results in significant reduction of Rb cell proliferation, migration, invasion, colony forming ability and in vivo tumor forming capability, while significantly increasing the rate of apoptosis of Rb cells.

STRING analysis further supported a functional association between UNC5D and NTN1 (P = 0.00392) (Fig. 6B). Finally, reciprocal co-immunoprecipitation assays confirmed a physical interaction between endogenous UNC5D and NTN1 in both Y79 and Weri-Rb1 cells (Fig. 6C, D). Together, these findings demonstrate that UNC5D associates with NTN1 in retinoblastoma cells and support the existence of a UNC5D–NTN1 signaling axis that may contribute to retinoblastoma progression.

## 4. Discussion

Retinoblastoma (Rb) remains the most common pediatric intraocular malignancy, and although RB1 loss is the initiating genetic event, additional molecular alterations are required for tumor progression [2,21]. In the present study, we identify UNC5D as a previously unrecognized oncogenic regulator in Rb and demonstrate that its depletion suppresses proliferation, migration, invasion, and tumor growth while promoting apoptosis in both in vitro and in vivo models. Mechanistically, our findings suggest that UNC5D contributes to PI3K/AKT pathway activation and associates with NTN1-mediated signaling in retinoblastoma cells.

UNC5D has traditionally been regarded as a dependence receptor with tumor-suppressive functions in multiple malignancies, including renal, prostate, thyroid, breast, and colorectal cancers [12–16,23]. In these tumor types, reduced UNC5D expression is frequently associated with promoter hypermethylation, loss of heterozygosity, or impaired apoptotic signaling [12–16]. In contrast, our findings demonstrate that UNC5D is highly expressed in retinoblastoma cells and promotes malignant phenotypes. These observations highlight the context-dependent nature of UNC5D signaling and suggest that its biological function may vary according to cellular lineage, ligand availability, and downstream signaling networks. Given the developmental origin of retinoblastoma from cone precursor-like cells, UNC5D may engage distinct signaling programs that favor cell survival and proliferation rather than dependence receptor-mediated apoptosis.

Transcriptomic profiling identified significant suppression of PI3K/AKT signaling following UNC5D knockdown. This pathway is a well-established regulator of cell growth, survival, and therapeutic resistance and has been implicated in retinoblastoma progression through genetic, epigenetic, and miRNA-mediated mechanisms [24–30]. Although total PIK3CA and AKT1 protein levels remained largely unchanged, phosphorylation of both PI3K and AKT was markedly reduced following UNC5D silencing in vitro and in xenograft tumors. These findings indicate that UNC5D primarily influences pathway activation rather than protein abundance. The observed discordance between AKT1 transcript and protein levels further suggests the involvement of post-transcriptional or post-translational regulatory mechanisms, which have been reported in several cancer models [31].

A major finding of this study is the identification of NTN1 as a potential mediator of UNC5D-associated signaling in retinoblastoma. NTN1 is a laminin-related guidance molecule that regulates cell survival through interactions with dependence receptors of the UNC5 family [32,33]. In the absence of NTN1, UNC5 receptors undergo caspase-mediated cleavage and trigger apoptotic signaling, whereas ligand engagement promotes cell survival [9]. Consistent with this model, NTN1 expression was elevated in retinoblastoma cells and decreased following UNC5D knockdown. Correlation analysis, molecular docking, co-immunoprecipitation, and expression studies collectively supported a close association between UNC5D and NTN1. Furthermore, the reduction of NTN1 expression observed following UNC5D depletion suggests that UNC5D may contribute to the maintenance of NTN1-associated signaling networks in retinoblastoma cells.

Importantly, our data demonstrate a robust physical interaction between UNC5D and NTN1 in both Y79 and Weri-Rb1 cells. Given the established role of NTN1 in suppressing dependence receptor-mediated apoptosis and promoting pro-survival signaling in several cancers [34–37], it is plausible that UNC5D–NTN1 interactions contribute to the apoptotic resistance observed in retinoblastoma. Although the precise molecular mechanisms linking UNC5D–NTN1 interactions to PI3K/AKT activation require further investigation, our findings support the existence of a functional UNC5D–NTN1 signaling axis in Rb.

Several limitations should be acknowledged. First, validation in primary retinoblastoma specimens was not performed. Second, although our findings support a functional association between UNC5D, NTN1, and PI3K/AKT signaling, additional rescue experiments will be required to establish direct causality. Third, the therapeutic efficacy of targeting the UNC5D–NTN1 axis remains to be evaluated in orthotopic and patient-derived models.

In conclusion, this study identifies UNC5D as a novel oncogenic regulator in retinoblastoma and provides evidence that UNC5D-associated NTN1 signaling could contribute to PI3K/AKT pathway activation, tumor cell survival, and disease progression. These findings expand the current understanding of retinoblastoma biology and nominate the UNC5D–NTN1 signaling axis as a potential therapeutic target for future investigation.

## 5. Conclusion

Retinoblastoma arises from the convergence of multiple signaling alterations, underscoring the need to define novel oncogenic drivers. Here, we provide the first evidence that UNC5D functions as an oncogene in Rb, promoting proliferation, migration, and tumorigenesis while suppressing apoptosis in vitro and in vivo. Mechanistically, UNC5D sustains PI3K/AKT pathway activation and engages in a strong interaction with its ligand NTN1, thereby facilitating tumor progression. These findings not only advance understanding of Rb biology but also nominate the UNC5D–NTN1–PI3K/AKT axis as a promising target for future diagnostic and therapeutic strategies aimed at improving outcomes for affected children.

## List of abbreviations

Rb: Retinoblastoma
NCBI-GEO: National Center for Biotechnology Information-Gene Expression Omnibus
UNC5D: Uncoordinated-5 Netrin Receptor D
RPMI-1640: Roswell Park Memorial Institute-1640
DMEM: F12 Dulbecco’s Modified Eagle Medium-Ham’s F-12
PBS: Phosphate Buffered Saline
FBS: Fetal Bovine Serum
shRNA: short hairpin RNA
cDNAs: complementary DNAs
SDS-PAGE: Sodium Dodecyl Sulfate Polyacrylamide Gel Electrophoresis
OCT: Optimal Cutting Temperature compound
HRP/DAB (ABC): Horseradish Peroxidase /3,3’-Diaminobenzidine (Avidin-Biotin Complex)
RIPA: Radio-immunoprecipitation assay
NTN1: Netrin 1
DEG: Differentially Expressed gene
STAR: Spliced Transcripts Alignment to a Reference
GO: Gene ontology
NES: Normalized Enrichment Score
Bcl-2: B-cell lymphoma 2
PI3K: Phosphoinositide 3-kinase
pPI3K: phospho Phosphoinositide 3-kinase
AKT: Ak strain transforming
pAKT: phosphor-Ak strain transforming
RCSB PDB: Research Collaboratory for Structural Bioinformatics Protein Data Bank
TCGA: The Cancer Genome Atlas
GEPIA: Gene Expression Profiling Interactive Analysis
UALCAN: The University of Alabama at Birmingham Cancer data analysis portal
H&E staining: Haematoxylin & Eosin staining.

## Acknowledgements

A.K. acknowledges financial support from the University Grants Commission (UGC), India (Award No. 16-[Dec-2017/2018(NET/CSIR)]). The authors thank Ms. Chirasmita Das, Mr. Md. Maqsood Ahamad Khan, Dr. Sandeepan Das, Mrs. Ruby Banerjee, and Mrs. Payel Karmakar for their technical assistance. S.K.G. gratefully acknowledges the Director, CSIR–Indian Institute of Chemical Biology (CSIR-IICB), Kolkata, for continuous support and encouragement.

## Declarations

### Author Contributions

A.K. conceived and performed the experiments, analyzed data, and drafted the manuscript. S.B. performed animal experiments and contributed to data analysis. Sh.B. conducted bioinformatic analyses and contributed to data interpretation. A.K.S. participated in study investigation, project administration, and manuscript review. S.K.G. conceived and supervised the study, secured funding, interpreted data, and revised the manuscript. All authors reviewed, approved, and agreed to the published version of the manuscript.

### Ethics Approval

All animal experiments were approved by the Institutional Animal Ethics Committee of CSIR–Indian Institute of Chemical Biology, Kolkata, India (Approval No. IICB/AEC/JAN/2024/12), and were conducted in accordance with institutional guidelines and national regulations governing the care and use of laboratory animals.

### Availability of data and materials

The RNA sequencing datasets generated during the current study have been deposited in the NCBI Gene Expression Omnibus (GEO) under accession number GSE306604. All other data supporting the findings of this study are available from the corresponding author upon reasonable request.

### Consent for publication

All the authors have evaluated the final version of the manuscript and given their approval for publication.

