## Supplementary File for "UNC5D Promotes Retinoblastoma Progression Through Association with NTN1 and Activation of PI3K/AKT Signaling"

**Table S1:**

**List of the gene specific shRNA sequences:**

| **Gene Name** | **Target Sequences (5ʹ-3ʹ)** |
| --- | --- |
| sh-UNC5D | ATGGG GAGAGCGGCG GCCACCGCAG GCGGCGGCGG AGGGGCGCGC CGCTGGCTCC CGTGGCTGGG GCTGTGCTTC TGGGCGGCAG GGACCGCGGC TGCCCGAGGA ACTGACAATG GCGAAGCCCT TCCCGAATCC ATCCCATCAG CTCCTGGGAC ACTGCCTCAT TTCATAGAGG AGCCAGATGA TGCTTATATT ATCAAGAGCA ACCCTATTGC ACTCAGGTGC AAAGCGAGGC CAGCCATGCA GATATTCTTC AAATGCAACG GCGAGTGGGT CCATCAGAAC GAGCACGTCT CTGAAGAGAC TCTGGACGAG AGCTCAGGTT TGAAGGTCCG CGAAGTGTTC ATCAATGTTA CTAGGCAACA GGTGGAGGAC TTCCATGGGC CCGAGGACTA TTGGTGCCAG TGTGTGGCGT GGAGCCACCT GGGTACCTCC AAGAGCAGGAAGGCCTCTGT GCGCATAGCC TATTTACGGA AAAACTTTGA ACAAGACCCA CAAGGAAGGG AAGTTCCCATTGAAGGCATG ATTGTACTGC  ACTGCCGCCC ACCAGAGGGA GTCCCTGCTG CCGAGGTGGA ATGGCTGAAA AATGAAGAGC CCATTGACTC TGAACAAGAC  GAGAACATTG ACACCAGGGC TGACCATAAC CTGATCATCA GGCAGGCACG GCTCTCGGAC TCAGGAAATT ACACCTGCAT  GGCAGCCAAC ATCGTGGCTA AGAGGAGAAG CCTGTCGGCC ACTGTTGTGG TCTACGTGAA TGGAGGCTGG TCTTCCTGGA  CAGAGTGGTC AGCCTGCAAT GTTCGCTGTG GTAGAGGATG GCAGAAACGT TCCCGGACCT GCACCAACCC AGCTCCTCTC  AATGGTGGGG CCTTTTGTGA GGGAATGTCA GTGCAGAAAA TAACCTGCAC TTCTCTTTGT CCTGTGGATG GGAGCTGGGA  AGTGTGGAGC GAATGGTCCG TCTGCAGTCC AGAGTGTGAA CATTTGCGGA TCCGGGAGTG CACAGCACCA CCCCCGAGAA  ATGGGGGCAA ATTCTGTGAA GGTCTAAGCC AGGAATCTGA AAACTGCACA GATGGTCTTT GCATCCTAGA TAAAAAACCT  CTTCATGAAA TAAAACCCCA AAGCATTGAG AATGCCAGCG ACATTGCTTT GTACTCGGGC TTGGGTGCTG CCGTCGTGGC  CGTTGCAGTC CTGGTCATTG GTGTCACCCT TTACAGACGG AGCCAGAGTG ACTATGGCGT GGACGTCATT GACTCTTCTG CATTGACAGG TGGCTTCCAG ACCTTCAACT TCAAAACAGT CCGTCAAGGT AACTCCCTGC TCCTGAATTC TGCCATGCAG  CCAGATCTGA CAGTGAGCCG GACATACAGC GGACCCATCT GTCTGCAGGA CCCTCTGGAC AAGGAGCTCA TGACAGAGTC  CTCACTCTTT AACCCTTTGT CGGACATCAA AGTGAAAGTC CAGAGCTCGT TCATGGTTTC CCTGGGAGTG TCTGAGAGAG  CTGAGTACCA CGGCAAGAAT CATTCCAGGA CTTTTCCCCA TGGAAACAAC CACAGCTTTA GTACAATGCA TCCCAGAAAT  AAAATGCCCT ACATCCAAAA TCTGTCATCA CTCCCCACAA GGACAGAACT GAGGACAACT GGTGTCTTTG GCCATTTAGG  GGGGCGCTTA GTAATGCCAA ATACAGGGGT GAGCTTACTC ATACCACACG GTGCCATCCC AGAGGAGAAT TCTTGGGAGA  TTTATATGTC CATCAACCAA GGTGAACCCA GCCTCCAGTC AGATGGCTCT GAGGTGCTCC TGAGTCCTGA AGTCACCTGT  GGTCCTCCAG ACATGATCGT CACCACTCCC TTTGCATTGA CCATCCCGCA CTGTGCAGAT GTCAGTTCTG AGCATTGGAA  TATCCATTTA AAGAAGAGGA CACAGCAGGG CAAATGGGAG GAAGTGATGT CAGTGGAAGA TGAATCTACA TCCTGTTACT  GCCTTTTGGA CCCCTTTGCG TGTCATGTGC TCCTGGACAG CTTTGGGACC TATGCGCTCA CTGGAGAGCC AATCACAGAC  TGTGCCGTGA AGCAACTGAA GGTGGCGGTT TTTGGCTGCA TGTCCTGTAA CTCCCTGGAT TACAACTTGA GAGTTTACTG  TGTGGACAAT ACCCCTTGTG CATTTCAGGA AGTGGTTTCA GATGAAAGGC ATCAAGGTGG ACAGCTCCTG GAAGAACCAA  AATTGCTGCA TTTCAAAGGG AATACCTTTA GTCTTCAGAT TTCTGTCCTT GATATTCCCC CATTCCTCTG GAGAATTAAA  CCATTCACTG CCTGCCAGGA AGTCCCGTTC TCCCGCGTGT GGTGCAGTAA CCGGCAGCCC CTGCACTGTG CCTTCTCCCT  GGAGCGTTAT ACGCCCACTA CCACCCAGCT GTCCTGCAAA ATCTGCATTC GGCAGCTCAA AGGCCATGAA CAGATCCTCC  AAGTGCAGAC ATCAATCCTA GAGAGTGAAC GAGAAACCAT CACTTTCTTC GCACAAGAGG ACAGCACTTT CCCTGCACAG  ACTGGCCCCA AAGCCTTCAA AATTCCCTAC TCCATCAGAC AGCGGATTTG TGCTACATTT GATACCCCCA ATGCCAAAGG  CAAGGACTGG CAGATGTTAG CACAGAAAAA CAGCATCAAC AGGAATTTAT CTTATTTCGC TACACAAAGT AGCCCATCTG  CTGTCATTTT GAACCTGTGG GAAGCTCGTC ATCAGCATGA TGGTGATCTT GACTCCCTGG CCTGTGCCCT TGAAGAGATT  GGGAGGACAC ACACGAAACT CTCAAACATT TCAGAATCCC AGCTTGATGA AGCCGACTTC AACTACAGCA GGCAAAATGG  ACTCTAG |
| sh-NC | CCTAAGGTTAAGTCGCCCTCGCTCGAGCGAGGGCGACTTAACCTTAGG |

**Table S2:**

**List of the oligonucleotide primer sequences of the selected genes employed in the qRT-PCR experiment:**

| **Name of the gene** | **Forward Primer sequence** | **Reverse Primer sequence** |
| --- | --- | --- |
| UNC5D | 5ʹ-ATTCGACTCGGGACCCTCAT-3ʹ | 5ʹ-ATTGTCAGTTCCTCGGGCAG-3ʹ |
| NTN1 | 5ʹ-TGACGGGTATCACCTGCAAC-3ʹ | 5ʹ-AGGAATCGCAGTCTTCAGGC-3ʹ |
| PIK3CA | 5ʹ-AGAGCCCCGAGCGTTTC-3ʹ | 5ʹ-TCACCTGATGATGGTCGTGG-3ʹ |
| AKT1 | 5ʹ-CTGCACAAACGAGGGGAGTA-3ʹ | 5ʹ-GCGCCACAGAGAAGTTGTTG-3ʹ |
| GAPDH | 5ʹ-TCGGAGTCAACGGATTTGGT-3ʹ | 5ʹ-TTCCCGTTCTCAGCCTTGAC-3ʹ |

**Table S3:**

**List of the antibodies used in western blot, immunofluorescence, immunohistochemistry and co-immunoprecipitation experiments:**

| **Name of the antibodies** | **Catalogue No.** | **Concentration & Application** |
| --- | --- | --- |
| Anti-UNC5D/UNC5H4 | BS-11494R, Bioss | 1:1000 (WB)  (Co-IP) |
| Anti-cyclin D1 | 2978T, CST | 1:1000 (WB) |
| Anti-cleaved caspase 3(p12) | A19664,ABclonal | 1:1000 (WB) |
| Anti-Bcl2(D17C4) | 3498T, CST | 1:1000 (WB) |
| Anti- PCNA(D3H8P) | 13110T, CST | 1:100 (IHC) |
| Anti-PI3K(p110 α) | A22730, ABclonal | 1:3000 (WB) |
| Anti-AKT(C67E7) | 4691S, CST | 1:1000 (WB) |
| Anti-pPI3K(PI3KP85α/γ/β-  Y467/Y199/Y464) | AP0854, ABclonal | 1:1000 (WB) |
| Anti-pAKT(S473-D9E) | 4060S, CST | 1:1000 (WB) |
| Anti- NTN1 | A16236, ABclonal | 1:1000 (WB)  (Co-IP) |
| Anti- β actin | BS-0061R, Bioss | 1:1000 (WB) |
| HRP conjugated goat anti-rabbit secondary antibody | 7074S, CST | 1:3000 (WB) |
| Normal rabbit IgG | sc-2027, Santa-Cruz Biotechnology, USA | (Co-IP) |
| Alexa Fluor 568 goat anti-rabbit IgG (H+L) | Invitrogen, A11011 | 1:400 (Immunofluorescence) |

**Supplementary method 1.**

**Single-cell RNA sequencing analysis**

The single-cell RNA-seq dataset GSE249995, comprising two intraocular and two extraocular retinoblastoma (RB) samples, was obtained from the Gene Expression Omnibus (GEO) [1]. Raw count data (barcode, feature, and matrix files) were processed in R (v4.5.1) using the Seurat package. Cells were filtered by unique feature counts and mitochondrial gene percentage, followed by normalization (NormalizeData) and scaling (ScaleData). Highly variable genes were identified with FindVariableFeatures. Dimensionality reduction was performed by PCA, UMAP (RunUMAP), and t-SNE (RunTSNE). Cell-type annotation was based on canonical markers reported by Liu et al. [1]. UNC5D expression across cell types was visualized using violin plots.

**Supplementary method 2.**

**Cell culture**

ShSY-5Y (CRL-2266), Hela (CCL-2) and HEK-293T (CRL-3216) cells were purchased from ATCC and Huh7 (CVCL-0336) was purchased from Cellosaurus. Hela cells were cultured in RPMI-1640 medium (Gibco, Thermo Fisher Scientific) supplemented with 10% fetal bovine serum (FBS) (Gibco, Thermo Fisher Scientific) and 1% penicillin/streptomycin (Gibco, Thermo Fisher Scientific). ShSY-5Y, Huh7 and HEK-293T cells were maintained in ATCC-formulated DMEM/F12 supplemented with 10% FBS and 1% penicillin/streptomycin. Cultured cells were routinely maintained in an incubator at 37℃ within a humidified environment of 5% CO_2_. Cells were subcultured and the culture media were replaced every two days after attaining 80% confluency. Cultured cells were routinely tested for mycoplasma detection.

**Supplementary method 3.**

**Detailed protocol for RNA Sequencing**

Total RNA was extracted from both transfected (shNC and shUNC5D) and wild-type Y79 cells using RNeasy mini kit (74104, Qiagen) and kept frozen at -80℃ before proceeding for RNA sequencing. Quality assessment of the extracted RNAs was done by using RNA Screen Tapes (5067-5576, Agilent) and accurate quantitation was performed with the help of Qubit RNA BR Assay (Q10211, Invitrogen). A modified NEBNext RNA Ultra II (NEB-E7775) directional protocol is used to generate the libraries for RNA sequencing and index codes were added to each sample to assign sequences. Library quality assessment was done on the Agilent 5400 system (Agilent, USA). Pooling and sequencing of the qualified libraries were done on the Illumina NovaSeq 6000 using 2 × 150 paired-end reads with 40million reads/sample. Fastq- mcf Program (v1.05) and cutadapt (v2.5) were applied to trim the adapter sequences and low-quality reads from the raw FASTQ files. Contamination removal was done by Bowtie2 (v2.5.1). All the paired-end reads were aligned to the reference Human genome obtained from Ensembl database (GRCh38/hg38). [STAR](https://github.com/alexdobin/STAR) (2.7.3a) was used for alignment. For estimating the expressions of genes, the raw read counts were assessed using [featureCounts](https://www.ncbi.nlm.nih.gov/pubmed/24227677) (v2.0.0). Read count data were normalized using an R Bioconductor package, [DESeq2](http://bioconductor.org/packages/release/bioc/html/DESeq2.html). Furthermore, cufflinks (v2.2.1) was utilized to estimate gene expression based on the aligned reads. For every gene, the expression values were represented in fragments per kilobase per million (FPKM) units. Hierarchical clustering analysis for normalized counts was achieved using R (v3.5.2) and additional packages viz. ggplot2, ggrepel and reshape2. Then differential expression analysis was done using [DESeq2](http://bioconductor.org/packages/release/bioc/html/DESeq2.html). All the identified DEGs were sorted according to their corresponding p-values for further analysis. Gene Set Enrichment Analysis (GSEA) and pathway analysis were carried out for significantly differentially expressed up- and down-regulated protein-coding genes. [ShinyGO](https://david.ncifcrf.gov/) (v0.80) and Kyoto Encyclopedia of Genes and Genomes ([KEGG](https://www.genome.jp/kegg/)) were used for Gene Ontology (GO) and pathway analysis. GO analysis was performed within the biological process (BP) category where p<0.05 was deemed statistically significant. GSEA analysis was performed using an R Bioconductor package called [clusterProfiler](https://bioconductor.org/packages/release/bioc/html/clusterProfiler.html) (v3.14.3) to unravel the functional significance of the DEGs.

(A)

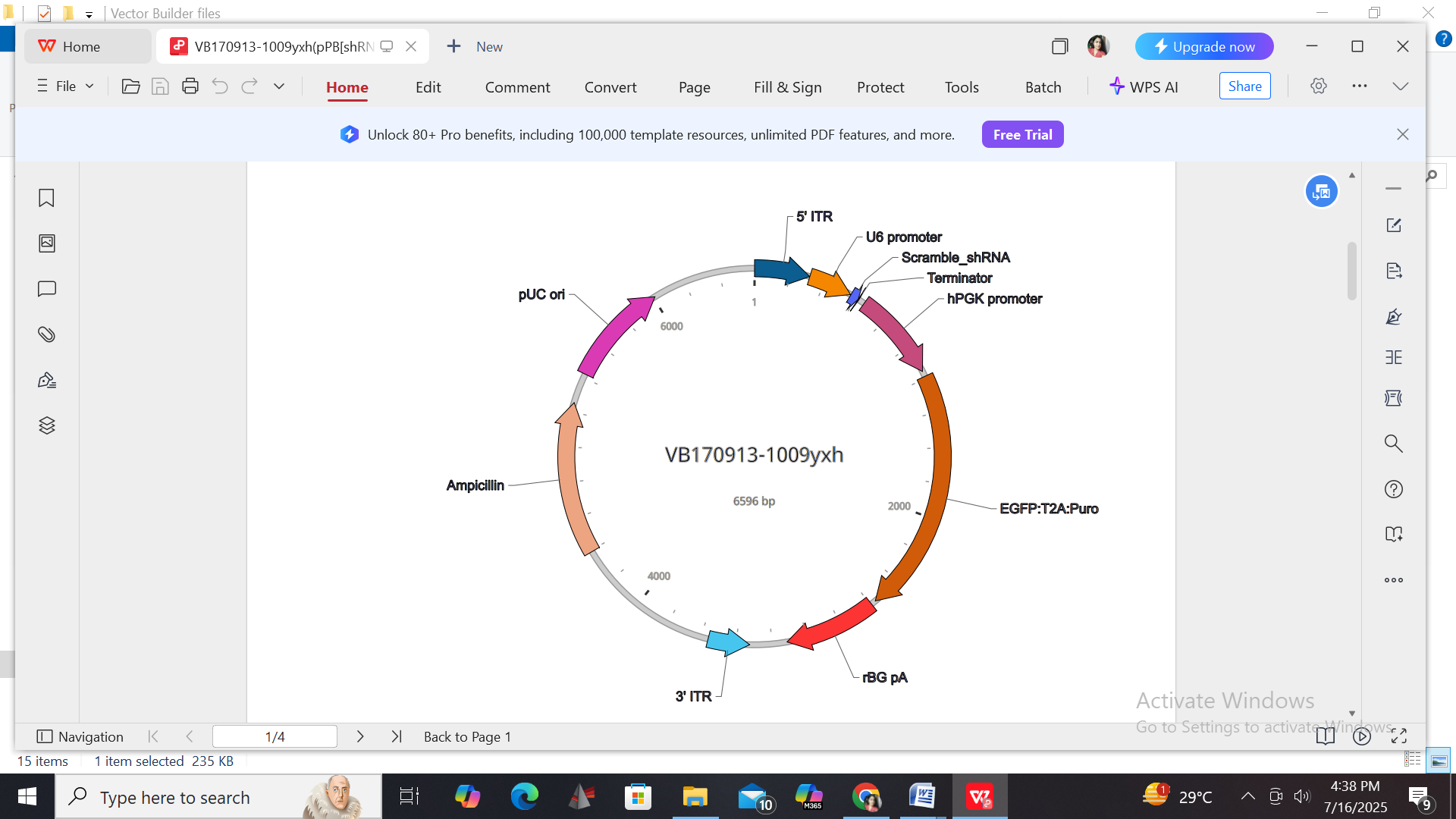

(B)

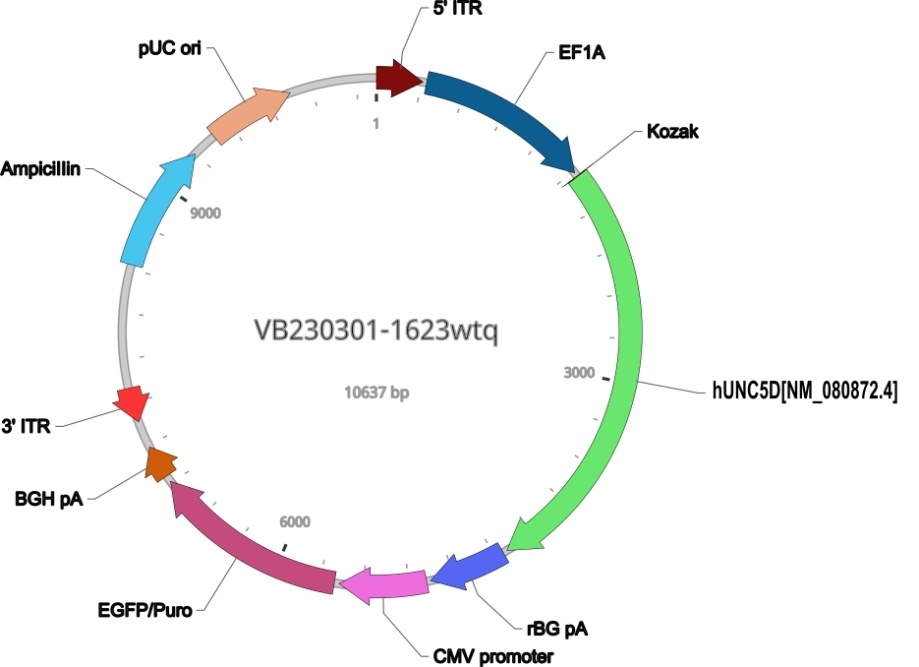

**Fig.S1** Vector maps of the respective shRNA constructs. (A) shNC (scramble_shRNA); (B) shUNC5D (shRNA targeting UNC5D).

(A)

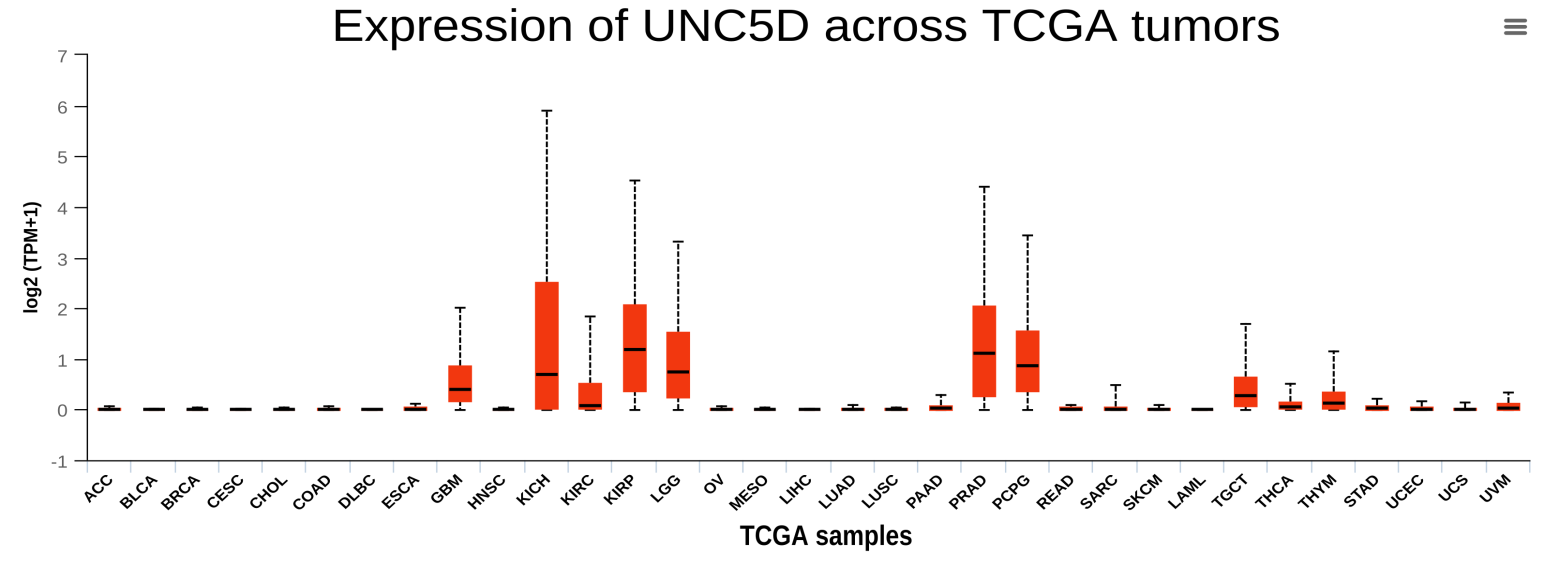

(B)

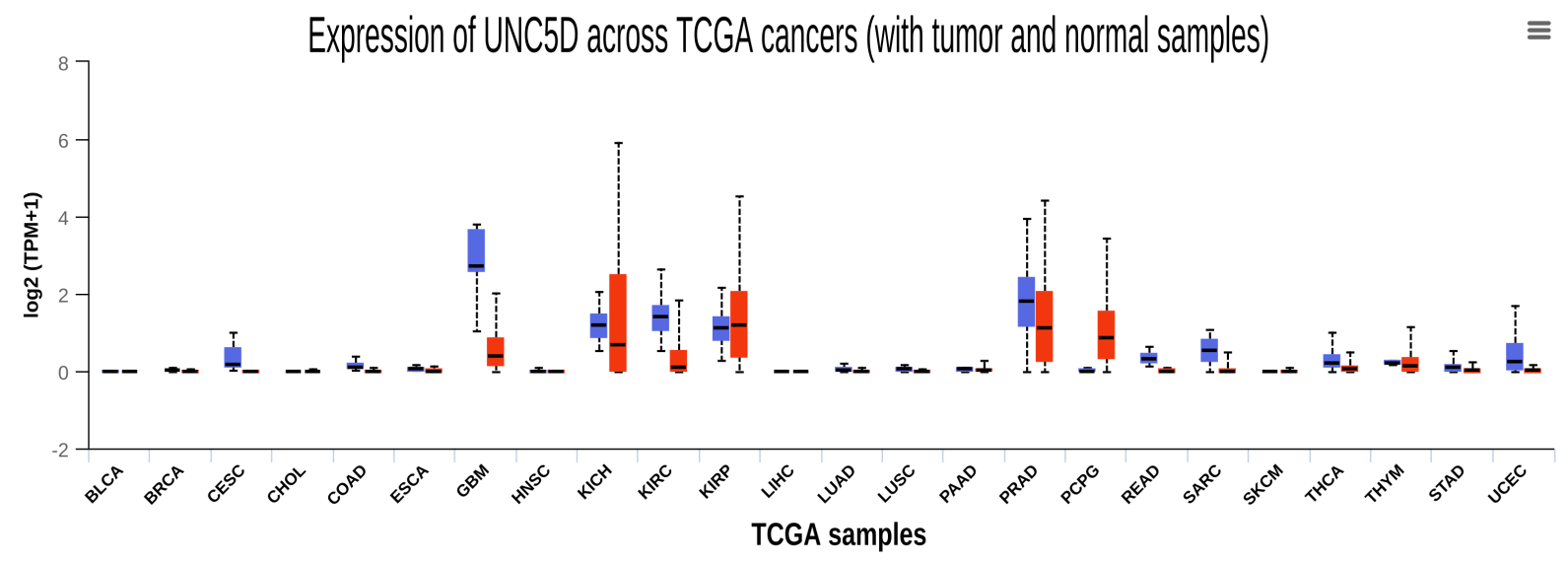

(C)

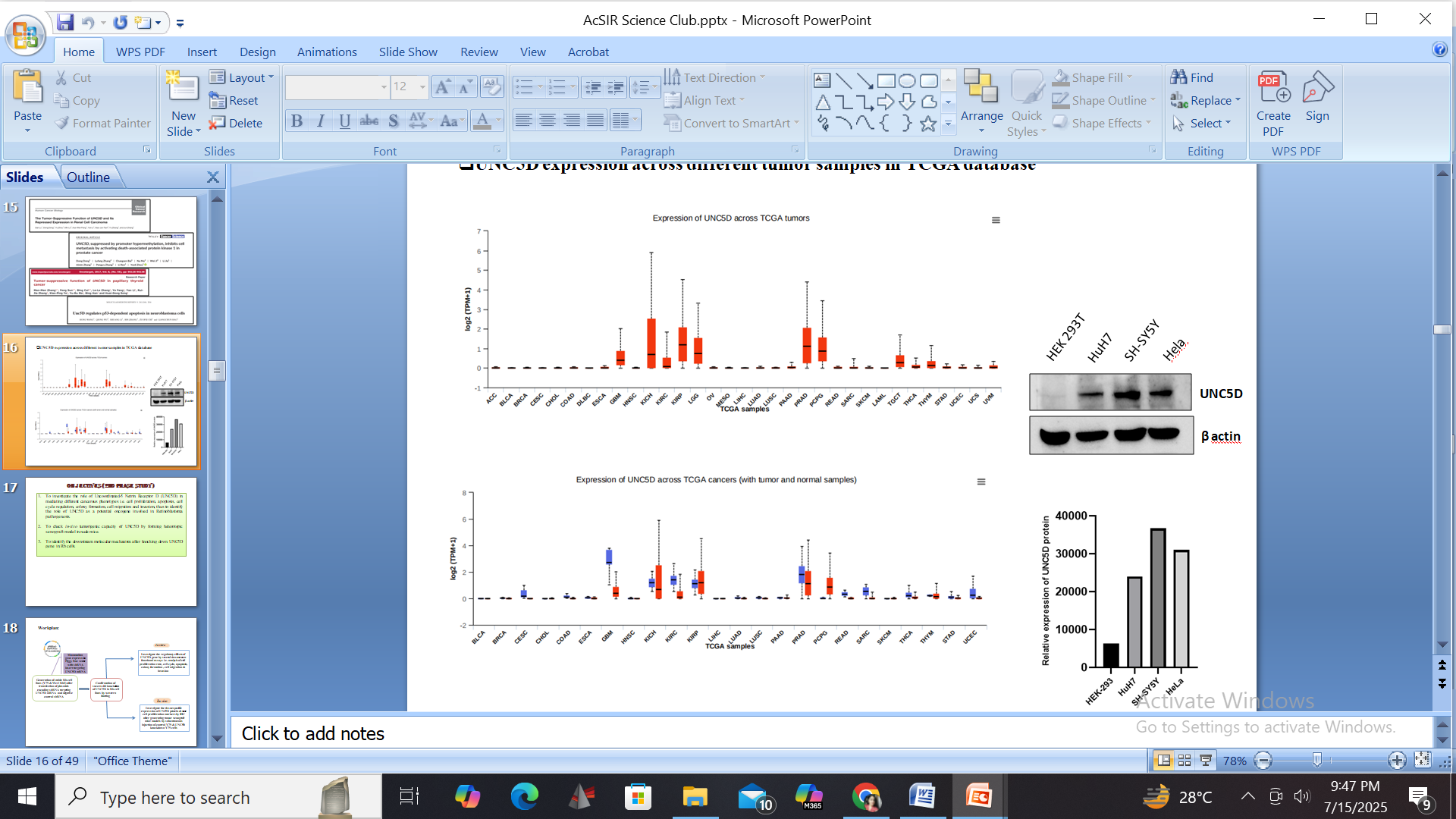

**Fig.S2. The expression of UNC5D is delineated through the analysis of publicly available TCGA datasets.** (A) UNC5D expression across different tumor samples in TCGA database. (B) Expression of UNC5D across TCGA cancers (including tumor and normal samples) as obtained from pan-cancer view analysis portal of The University of Alabama at Birmingham Cancer data analysis portal (UALCAN) server. (C) Relative protein expression of UNC5D in three different human cancer cell lines viz. Hela (cervical cancer cell line), SH-SY5Y (neuroblastoma cell line) and HuH7 (hepatocellular carcinoma cell line) and non-cancerous HEK-293T cell line. β-actin was used here as loading control.

(A)

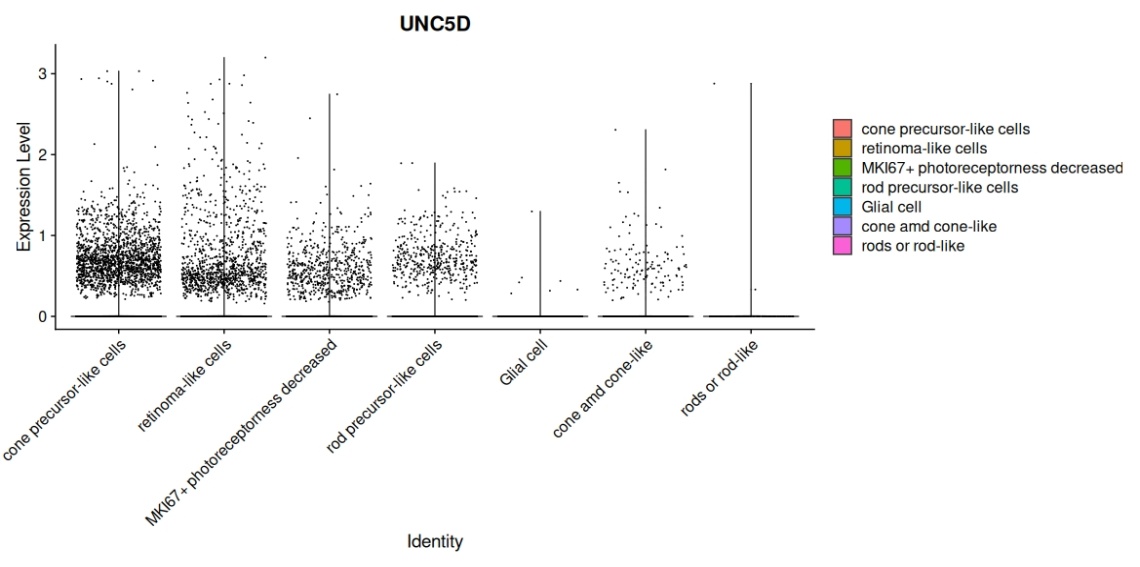

(B)

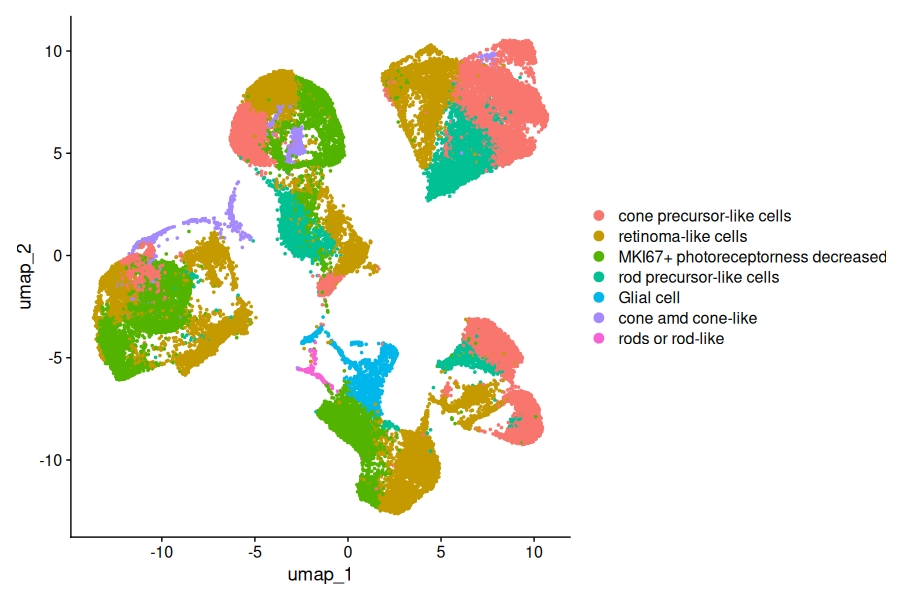

(C)

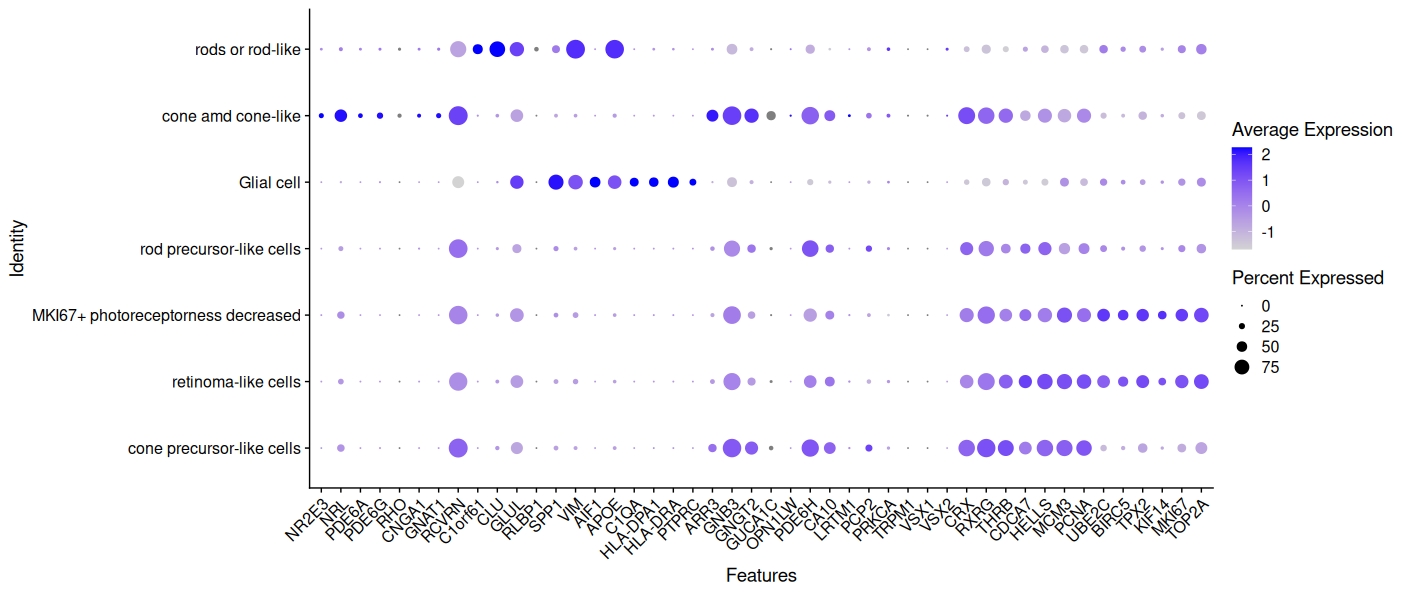

**Fig.S3.**(A) Violin plot showing the expression of UNC5D across different retinal cell types. (B) UMAP plot for all qualified cells. (C) Representative gene markers for each cell type in the retina. The scRNA-seq dataset 249995 was integrated and processed for this analysis.

(A)

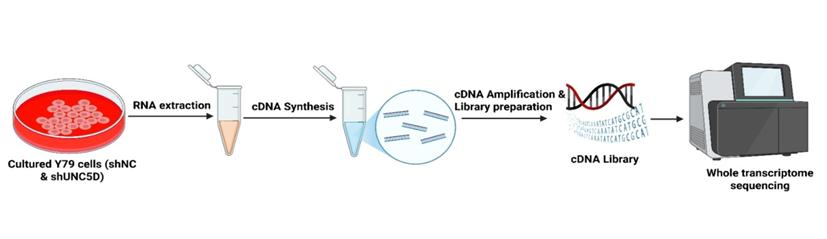

(B)

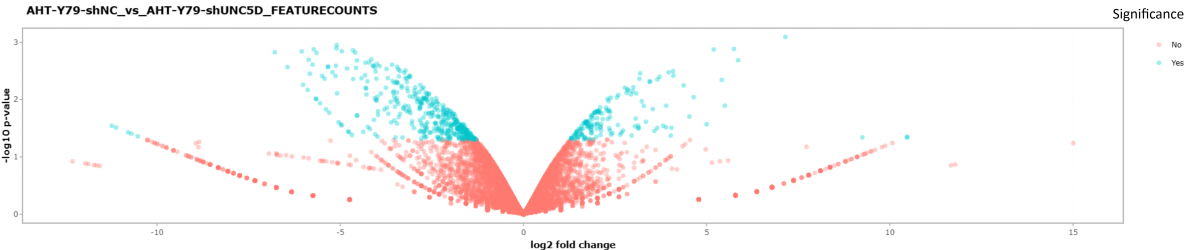

(C)

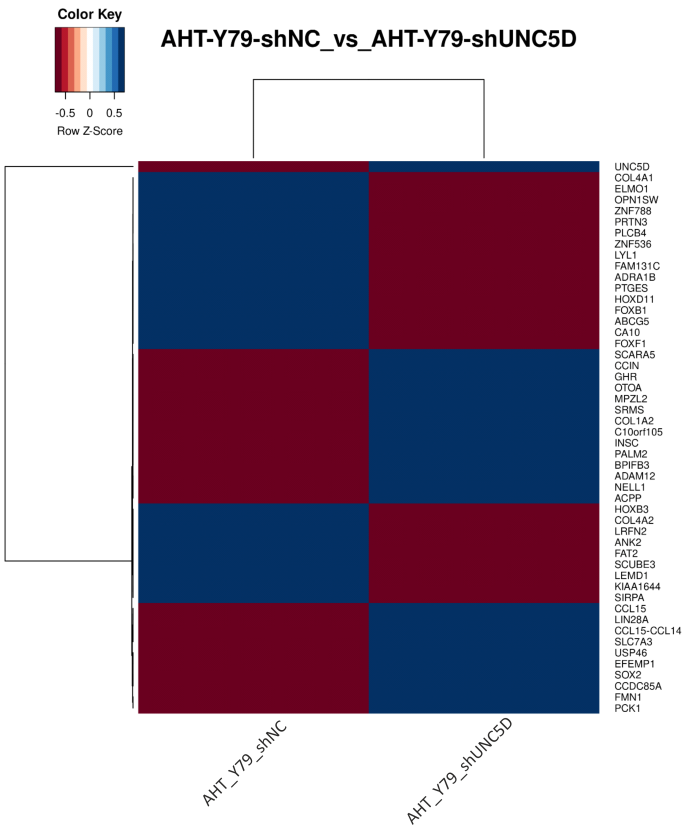
 (D)	**Upregulated**

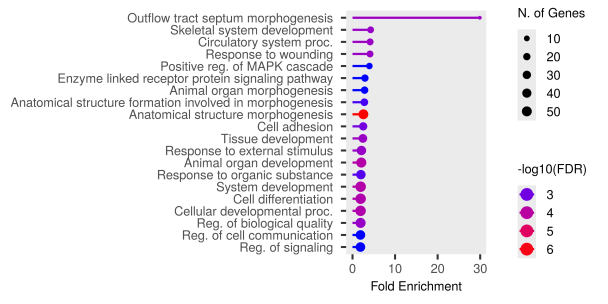

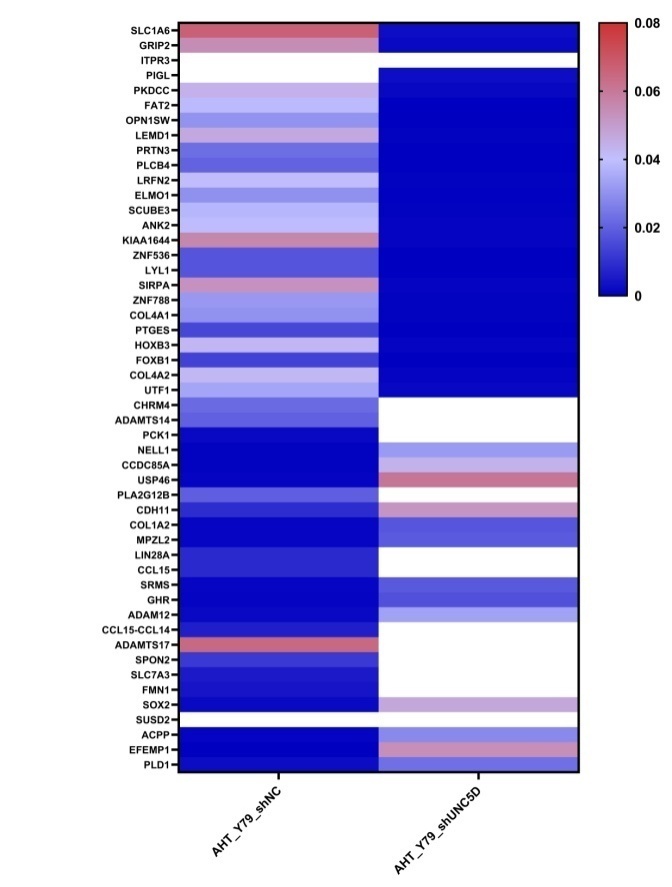

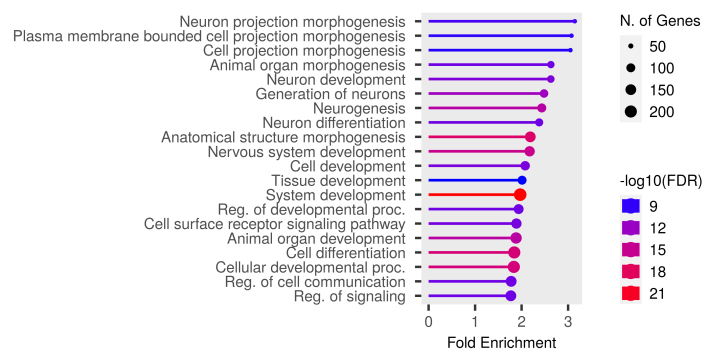

(E)	**Downregulated**

**Fig.S4. Whole transcriptome sequencing shows alteration in signaling pathway in response to UNC5D downregulation in retinoblastoma.** (A) Schematic of whole transcriptome sequencing in Y79 Rb cells after knocking down UNC5D gene as compared to the scramble control cells. (B) Volcano plot showing differential expression of protein coding genes (in shNC& shUNC5D groups). (C) Heatmap showing expression for top 25 and bottom 25 significantly differentially expressed protein coding genes (in shNC& shUNC5D groups). (D) Pathway enrichment analysis showing GO Biological process of upregulated DEGs [p<0.05, log_2_FC>+-2]. (E) Pathway enrichment analysis showing GO Biological process of downregulated DEGs [p<0.05, log_2_FC>+-2].

**Upregulated	Downregulated**

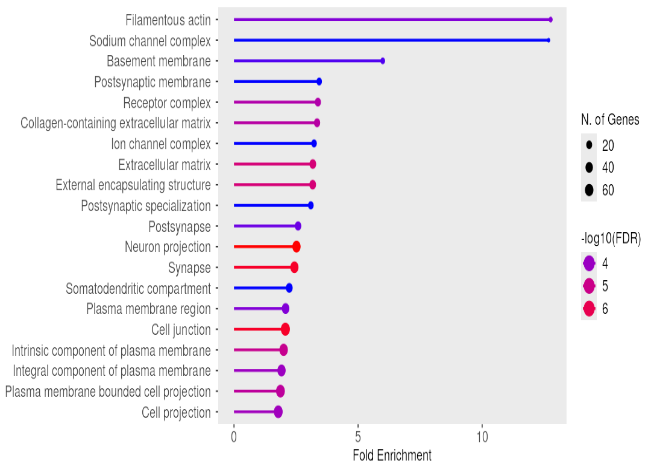

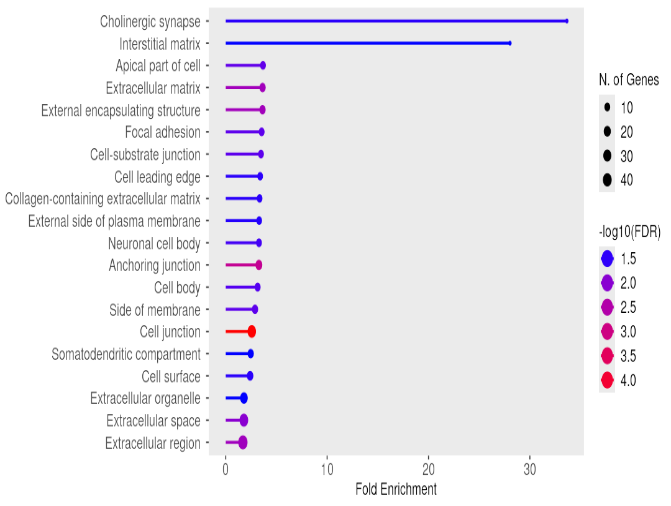

(B)

(A)

(C)	**Upregulated**

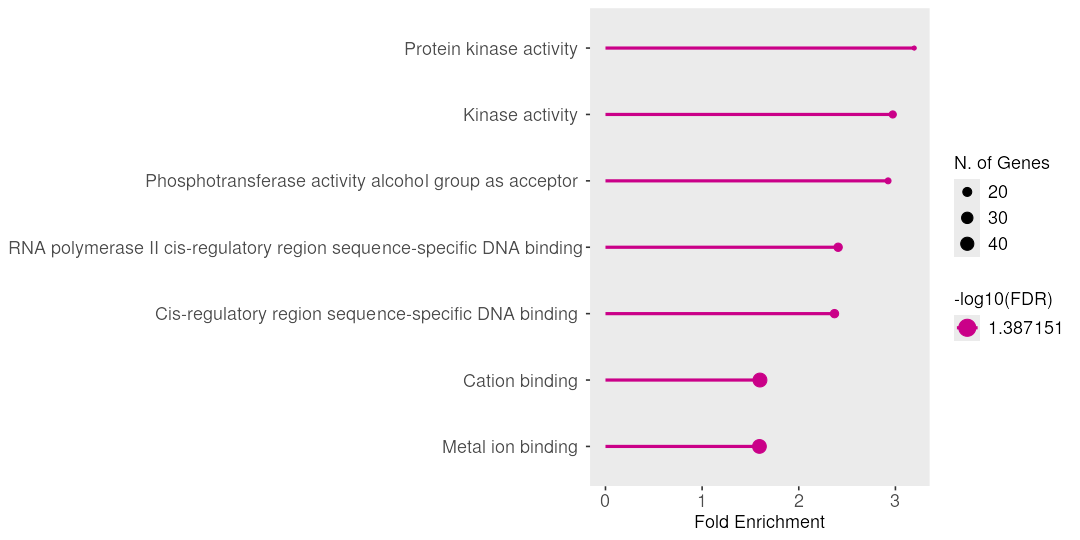

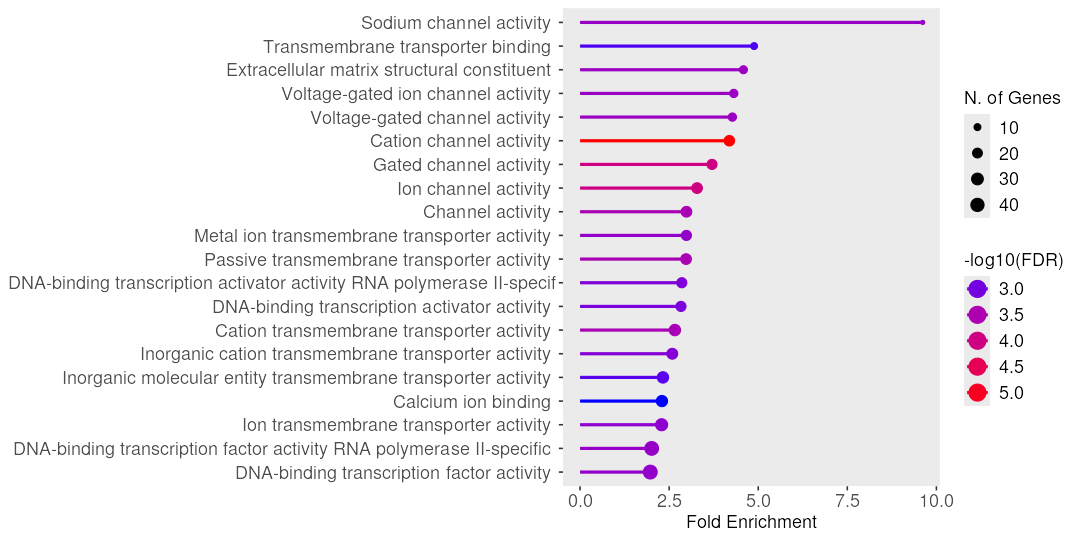
(D)	**Downregulated**

**Fig.S5.** (A & B) Pathway enrichment analysis showing GO Cellular component of upregulated & downregulated DEGs [p<0.05, log_2_FC>+-2]. (C & D) Pathway enrichment analysis showing GO Molecular function of upregulated & downregulated DEGs [p<0.05, log_2_FC>+-2].

(A)

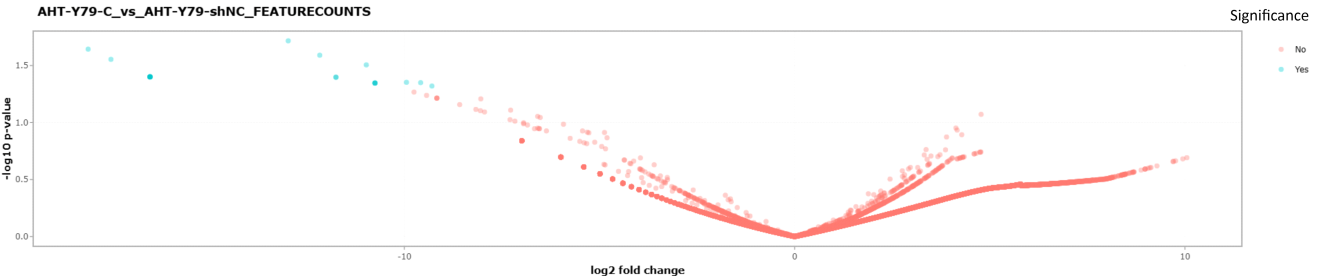

(B)

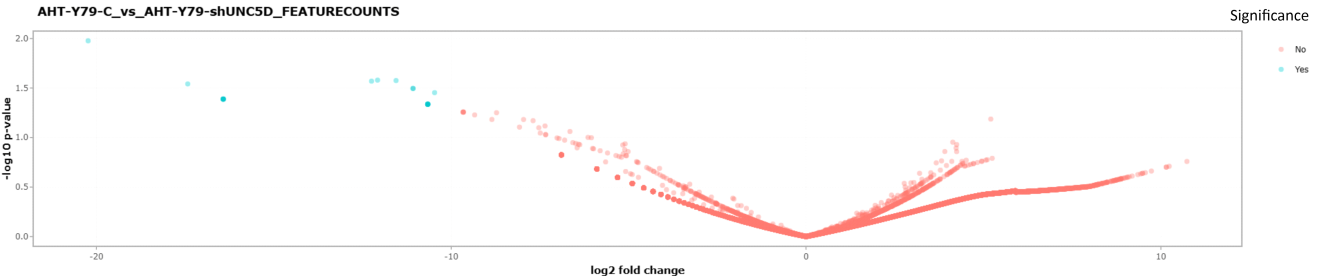

(C)	(D)

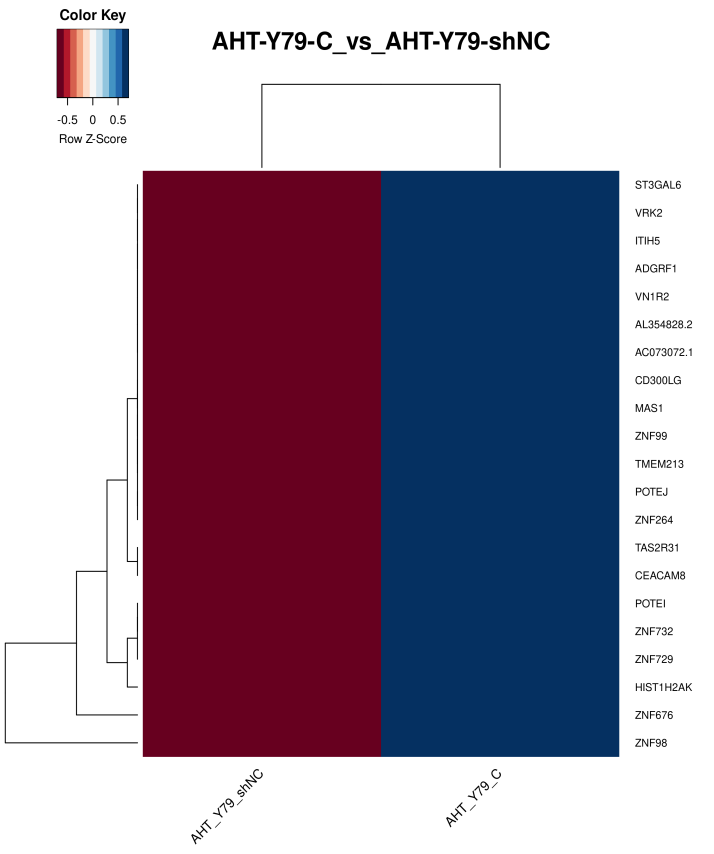

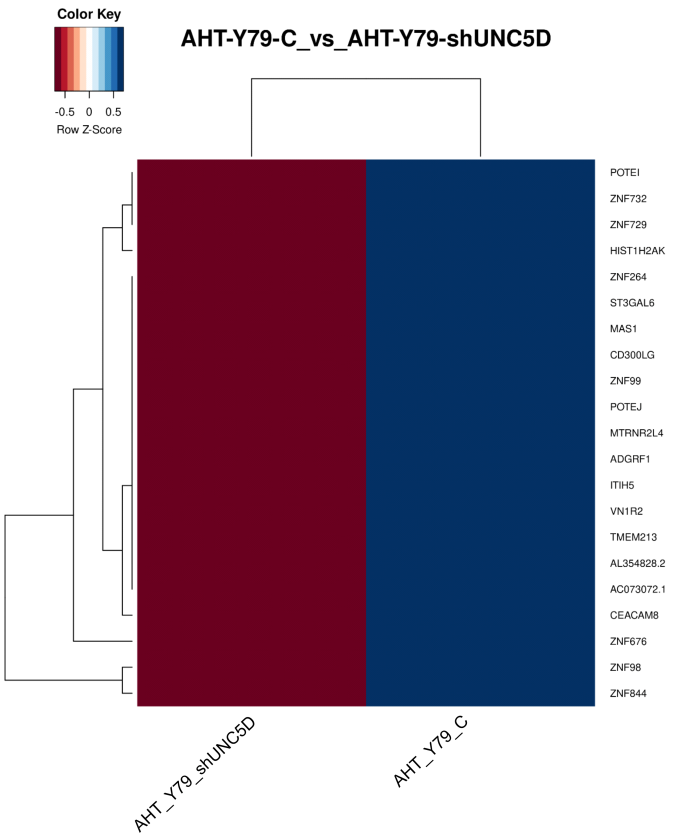

**Fig.S6.** (A) Volcano plot showing differential expression of protein coding genes (in wild type C & shNC groups). (B) Volcano plot showing differential expression of protein coding genes (in wild type C & shUNC5D groups). (C) Heatmap showing expression for top 21 significantly differentially expressed protein coding genes (in wild type C & shNC groups). (D) Heatmap showing expression for top 21 significantly differentially expressed protein coding genes (in wild type C & shUNC5D groups). [p<0.05, log_2_FC>+-2].

(A) (B)

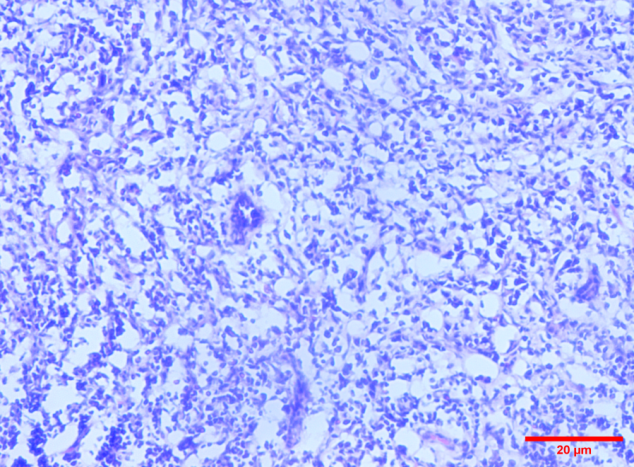

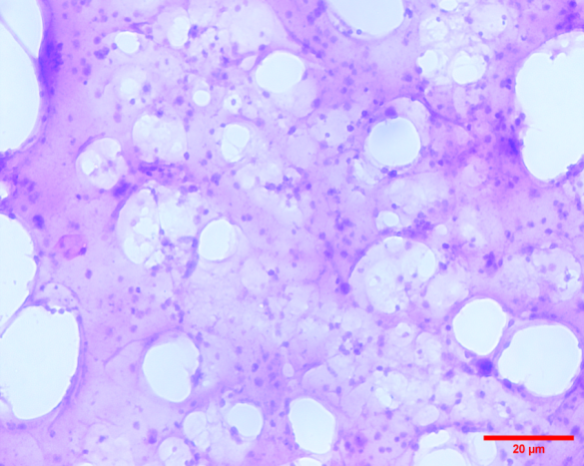

**shNC	shUNC5D**

**Fig.S7.** Representative H&E staining images of the xenograft tumor tissues. (A) shNC group; (B) shUNC5D group. (Magnification, 20×) (Scale bars, 20 μm).

(A)

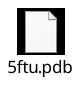

(B)

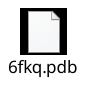

#### Fig.S8. Original PDB files of the above mentioned proteins used for the docking study. (A) Legacy PDB format of UNC5D (Tetrameric complex of Latrophilin 3, Unc5D and FLRT2); (B) Legacy PDB format of NTN1 (Thecrystalstructure ofafragmentof Netrin-1 incomplexwith afragmentofDraxin).
